# Early-life colonization by enterotoxigenic *Bacteroides fragilis* remodels gut epithelial stem cell states to drive colorectal cancer susceptibility

**DOI:** 10.64898/2026.09.28.755180

**Authors:** Seongmi K. Russell, Ezequiel Valguarnera, Jesse J. Pak, Juliane Bubeck Wardenburg

## Abstract

Early-life microbial exposures can shape lifelong disease risk, yet how developmental timing influences the host-microbe interaction remains unclear. During neonatal colonization by enterotoxigenic *Bacteroides fragilis* (ETBF), the metalloprotease toxin BFT enables lamina propria niche entry and remodeling of colonic epithelial cell states. Early-life ETBF exposure expands murine Lgr5^+^ stem cells, enhances Atoh1^+^ secretory differentiation, and induces distal colon-specific Wnt hyperactivation. These epithelial alterations coincide with increased formation of precancerous lesions in *Apc*^Min/+^ mice. ETBF preferentially forms intracellular bacterial aggregates within these lesions, establishing a persistent reservoir that reinforces epithelial remodeling. Colonization with non-toxigenic *B. fragilis* during this critical developmental window prevents lesion formation, revealing that developmental context dictates whether microbial colonization imprints protection or predisposition toward colorectal cancer.

## Main Text

The colonic microbiota is now recognized as a key determinant of host susceptibility to intestinal and extra-intestinal disease, fueling interest in strategies to modulate the microbiota to mitigate disease ^1–3^. Colorectal cancer (CRC) is the second leading cause of cancer-related deaths ^4,5^. In recent years, CRC incidence has increased among individuals under 50 years ^6,7^, suggesting that environmental risk factors, including acquisition of specific microbes, may exert a strong influence on disease susceptibility. Human studies reveal remarkable stability of the gut microbiome ^8–12^, indicating that early colonizers may imprint host metabolism, immunity, and long-term health. *Bacteroides fragilis* is among the earliest colonizers: while non-toxigenic *B. fragilis* (NTBF) strains promote mucosal immune tolerance ^13–16^, enterotoxigenic *B. fragilis* (ETBF) strains produce the zinc-dependent metalloprotease *B. fragilis* toxin (BFT) that elicits epithelial injury and colitis ^17,18^. ETBF is detected in up to 20% of healthy individuals ^19–23^, but is enriched in patients with inflammatory bowel disease ^19^, children with diarrhea or undernutrition ^24^, and is associated with human CRC lesions ^25,26^. ETBF accelerates tumorigenesis in *Apc*^Min/+^ mice ^21,27,28^, raising the possibility that this organism acts as an infectious carcinogen analogous to *Helicobacter pylori* ^3,26,29^. Unlike the conventional microbiota including NTBF which is restricted to the colonic lumen, ETBF exhibits unique niche acquisition behavior during early life. Using our neonatal colonization model, we recently showed that ETBF penetrates the inner mucus and lamina propria (LP) through goblet cell-associated antigen passages (GAPs) in a BFT-dependent manner during a restricted developmental window ^30,31^. As acquisition of this niche places BFT in proximity to the basolateral surface of colonic crypt stem cells, we hypothesized that toxin exposure may alter epithelial cell fate, remodel crypt dynamics, and thereby increase risk of colon tumorigenesis. To test this, we used a vertical transmission model in which ETBF-colonized pregnant dams enable natural colonization of offspring ^30^. This system avoids antibiotic-induced dysbiosis and permits longitudinal analysis of ETBF-host interactions. Using this approach, we define how early-life ETBF colonization reprograms epithelial development and drives precancerous transformation through BFT exposure.

### Fpn-dependent BFT activation during early-life ETBF colonization influences epithelial progenitor fate

We previously identified the cysteine protease Fragipain (Fpn) as the enzyme responsible for BFT activation ^32^. Prior work showed that Fpn is dispensable for BFT-mediated epithelial injury in adult intestinal infection, where colonic mucus-associated protease activity can mediate BFT cleavage, but is required for BFT activation and virulence in non-luminal bloodstream infection ^32^. To examine the role of Fpn-dependent BFT activation during early-life colonization, we used a vertical transmission model in which *B. fragilis*-colonized pregnant dams transmit bacteria to offspring during the neonatal period (Fig. 1A). Based on this compartment-specific requirement, we hypothesized that Fpn-dependent BFT activation may be important during early-life ETBF colonization after bacterial access to the LP, where exposure to luminal mucus-associated protease activity is limited. Consistent with this idea, neonatal colonization with ETBF Δ*fpn* resulted in reduced crypt elongation, producing an intermediate phenotype between wild-type ETBF (WT) and Δ*bft* strains (Fig. 1B, Extended Data Fig. 1A). Quantification at 2, 3, and 5 weeks of age confirmed that both BFT and Fpn are required for ETBF-induced crypt elongation, with Δ*fpn* displaying an intermediate effect (Fig. 1C, Extended Data Fig. 1B-C). Niche-specific CFU analysis showed that luminal colonization of Δ*fpn* was comparable to WT, but mucus and tissue colonization were significantly impaired by loss of Fpn (Fig. 1D). Δ*bft* colonization was reduced across all niches as expected (Fig. 1D). Confocal imaging of ETBF strains engineered to encode superfolder-GFP (sfGFP) ^30^ revealed diminished tissue colonization by ETBF Δ*fpn* compared to WT at 3 weeks of age (Fig. 1E). These results indicate that Fpn is required for local activation of BFT to promote efficient ETBF colonization of mucus and epithelial tissue during early life.

**Fig. 1.**
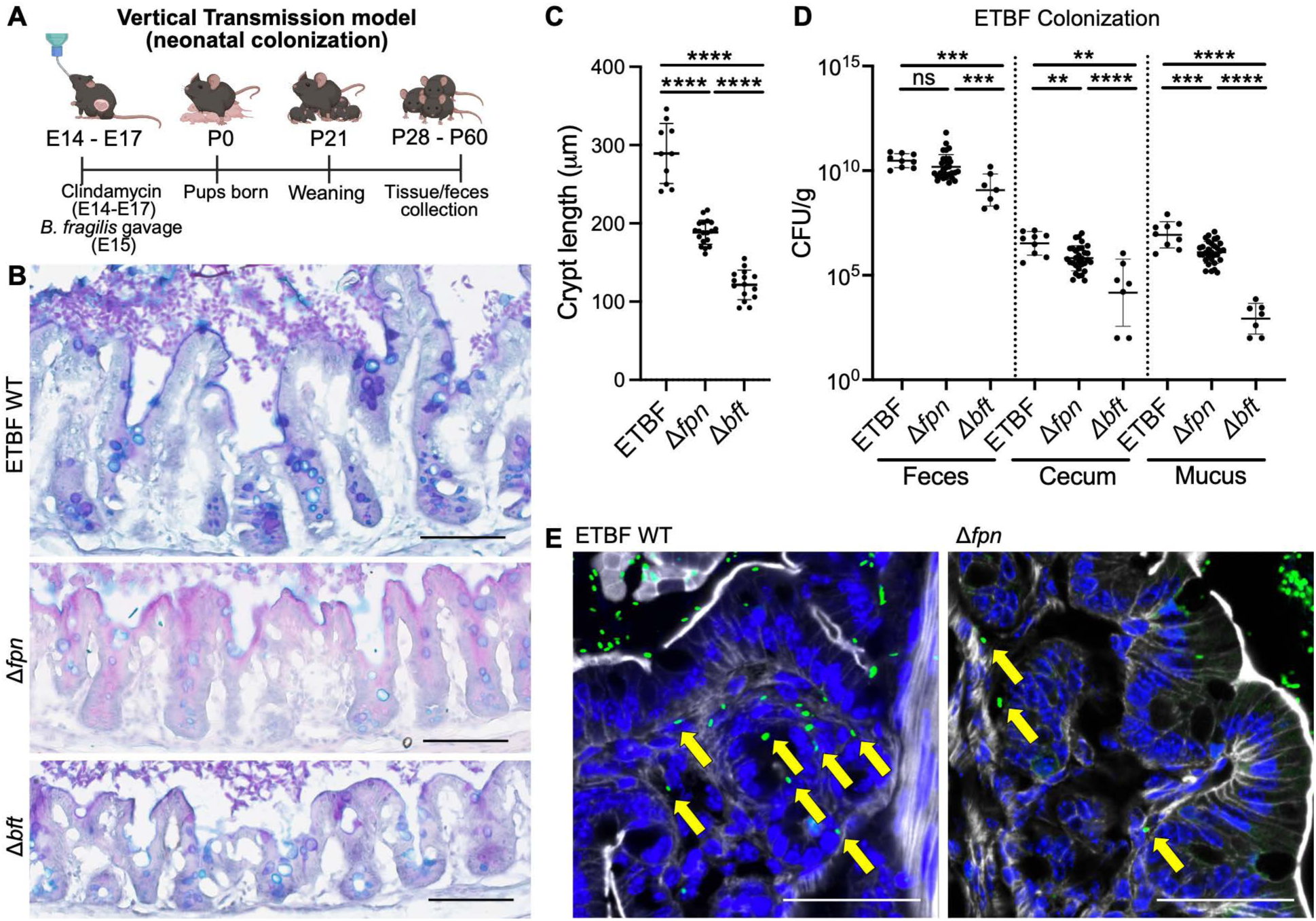
Early-life ETBF colonization promotes BFT-and Fpn-dependent tissue colonization and crypt remodeling. (A) Timeline of the vertical transmission model (neonatal colonization). Pregnant dams were treated with clindamycin at embryonic day 14-17 (E14-E17) and gavaged with *B. fragilis* at E15. Pups were analyzed at weaning (P21), and between P28-P60 for tissue and fecal samples. Created in BioRender. (B) Representative AB/PAS staining of cecal tissue from mice colonized with ETBF WT, Δ*fpn*, or Δ*bft* at 5 weeks of age. (C) Quantification of crypt length at 5 weeks of age. Crypt length was measured as the distance between the upper and lower ends of the crypt across multiple images. Data from 2-and 3-week colonization are shown in Extended Data Fig. 1. (D) ETBF WT, Δ*fpn*-, or Δ*bft*-colonized pups were analyzed at 3 weeks, and ETBF colonization (CFU/g) in feces, cecum, and mucus was quantified. (E) ETBF WT and Δ*fpn*-colonized ceca at 3 weeks were stained for F-actin (white) and DAPI (blue). Yellow arrows indicate intracellular ETBF (green) within the lamina propria. Scale bars, 100 μm. *p < 0.05, **p < 0.01, ***p < 0.001, ****p < 0.0001, ns = not significant.

To examine whether BFT is sufficient to alter epithelial proliferation, we utilized primary colonic organoids ^33^. Exposure of organoids to recombinant active BFT caused visible junctional disorganization and dense cellular centers within 3 hours (green and yellow arrows, Extended Data Fig. 2A, left), compared to organoids treated with inactive BFT (Extended Data Fig. 2B, left). Confocal imaging at 3 hours showed disrupted F-actin organization and surface discontinuity in BFT-treated organoids (Extended Data Fig. 2A, middle), whereas inactive BFT-treated organoids retained an intact epithelium (Extended Data Fig. 2B, middle). To determine whether these morphological changes were associated with epithelial proliferation, we assessed Ki67 expression. BFT-treated organoids exhibited increased Ki67 staining compared with inactive BFT-treated organoids (Extended Data Fig. 2, right), supporting increased proliferative activity following direct BFT exposure. These findings demonstrate that colonic epithelial cell exposure to BFT alone, in the absence of other microenvironmental stimuli, is sufficient to disrupt epithelial junctions and enhance proliferation, providing a strong rationale to investigate epithelial cell fate signaling changes during ETBF colonization.

Colonic epithelial homeostasis is maintained by Lgr5⁺ stem cells at the crypt base, which give rise to transit-amplifying cells that differentiate into absorptive or secretory lineages ^38–40^. Wnt signaling, most active at the crypt base, sustains stem cell maintenance and proliferation and is the principal pathway driving colorectal cancer (CRC) when mutationally activated ^41,42^. By contrast, Atoh1 directs secretory lineage commitment, including goblet cell differentiation ^43^. To investigate how BFT perturbs these pathways, we analyzed Lgr5 (Lgr5-EGFP-IRES-CreERT2), Wnt activity (TCF/Lef:H2B-GFP), and Atoh1 (R26-LSL-H2B-mCherry; Atoh1-Cre) reporter mice colonized with ETBF WT, Δ*fpn*, or Δ*bft* strains to visualize BFT-dependent mucosal remodeling at single-cell resolution in the cecum. Lgr5-eGFP mice revealed expansion of Lgr5⁺ intestinal stem cells along the crypt axis in ETBF WT-colonized colon, whereas Δ*fpn*, Δ*bft*, and uncolonized controls showed Lgr5^+^ cells confined to the crypt base (Fig. 2A, quantified adjacent to image). Although ETBF WT colonization expanded the Lgr5⁺ compartment, the fraction of Lgr5⁺ cells per crypt was unchanged, indicating that stem cell expansion was proportional to total crypt cell number (Extended Data Fig. 3A-B). Wnt activity reporter mice demonstrated enhanced Wnt activity localized to the crypt base in ETBF WT-colonized crypts, dependent on both Fpn and BFT (Fig. 2B, quantified adjacent to image; see also Extended Data Fig. 4). Atoh1 reporter mice showed expansion of Atoh1⁺ secretory progenitors in ETBF WT-colonized crypts compared with Δ*fpn*, Δ*bft*, and uncolonized controls (Fig. 2C, quantified adjacent to image), consistent with the observed increase in goblet cells upon ETBF colonization ^30^. Reporter specificity was validated using R26-LSL-H2B-mCherry; Atoh1-Cre^-/-^ littermate controls (Extended Data Fig. 5A-B). Epithelial mCherry⁺ cells were restricted to Cre^+/-^ tissues (Fig. 2C), whereas stromal background signal was observed in both Cre^+/-^ and Cre^-/-^ controls (Extended Data Fig. 5A), confirming that epithelial mCherry⁺ cells represent bona fide Atoh1⁺ secretory progenitor cells. Together, this data show coordinated induction of Lgr5, Wnt, and Atoh1 signaling, suggesting that BFT activation by Fpn within the colonic tissue is required to drive proliferation and promote secretory lineage differentiation during early-life ETBF colonization.

**Fig. 2.**
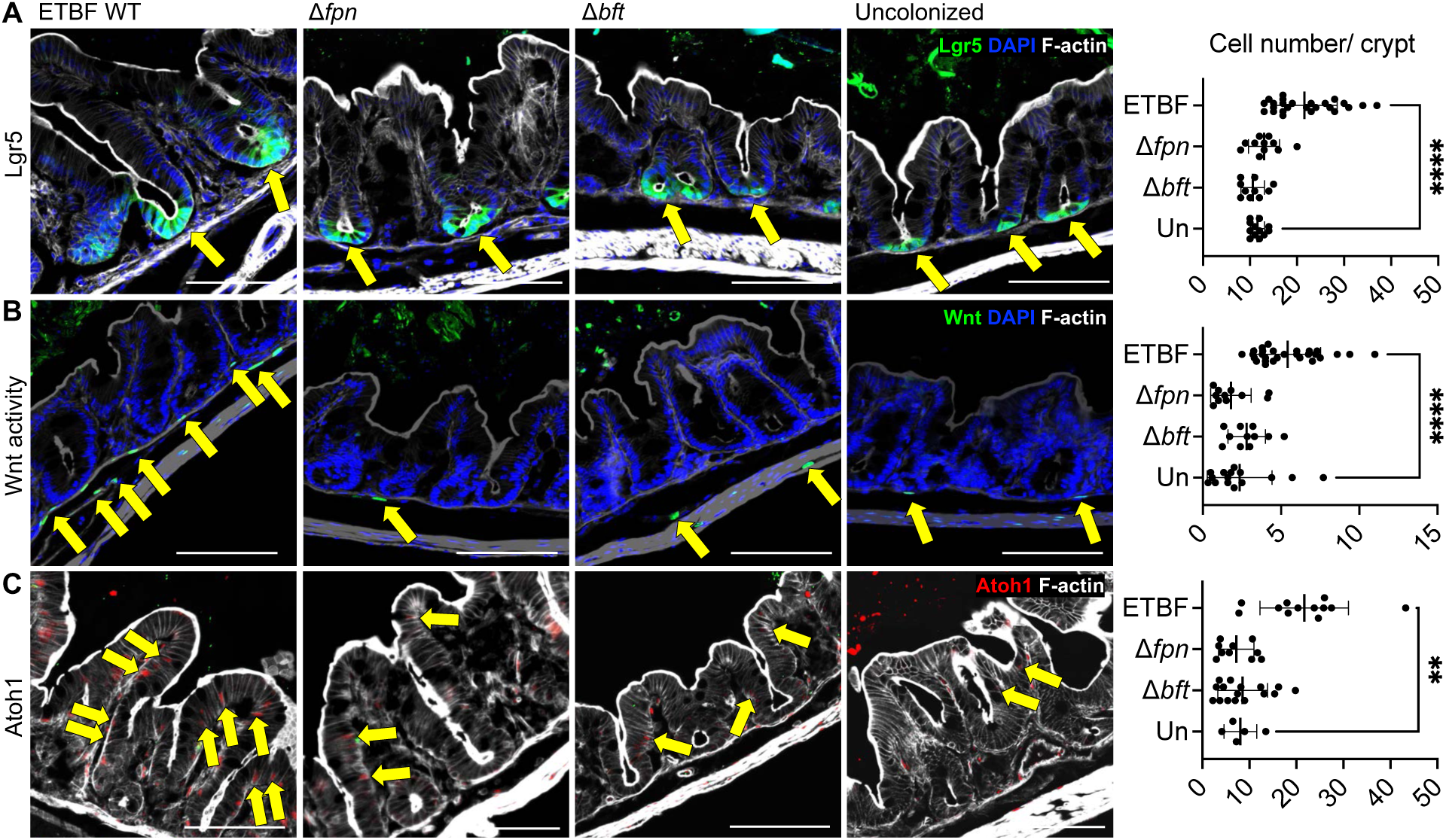
BFT and Fpn modulate *in vivo* cecal crypt cell fate. (A) Lgr5 (Lgr5-EGFP-IRES-CreERT2), (B) Wnt activity (TCF/Lef:H2B-GFP), and (C) Atoh1 (R26-LSL-H2B-mCherry; Atoh1-Cre) reporter mice were colonized with ETBF WT, Δ*fpn*, or Δ*bft* strains; uncolonized reporter mice served as controls. Yellow arrows indicate representative positive cells in each image. Quantification is shown adjacent to each image. Scale bars, 100 μm. **p < 0.01, ****p < 0.0001. Abbreviations: uncolonized (Un).

### ETBF colonization during early life initiates colorectal tumor precursors

We hypothesized that toxin-mediated tissue remodeling events may predispose susceptible hosts to colorectal tumorigenesis. To test whether early-life ETBF colonization promotes CRC initiation, we evaluated vertical transmission of ETBF in *Apc*^Min/+^ mice that harbor a point mutation in the adenomatous polyposis coli locus, leading to truncation of the Apc protein and promoting tumor development in heterozygous mutant mice ^44,45^. Pregnant *Apc*^Min/+^ dams were colonized with ETBF WT, Δ*fpn* or Δ*bft* strains, and colons from 4-week-old *Apc*^Min/+^ pups were analyzed (Fig. 3A-D). Aberrant crypt foci (ACF), the earliest CRC precursor lesions, were visualized by methylene blue (MB) staining and classified as primal (1-2 crypts, yellow), intermediate (3-6 crypts, blue), or advanced (>7 crypts, red), (Fig. 3A, lower panels, magnified views), as previously defined in rodent models of colon cancer ^46–49^ and also identified in the human colon ^50^. ETBF WT-colonized pups developed significantly more ACF compared to Δ*fpn*, Δ*bft*, or uncolonized controls (Fig. 3E-F). Notably, Δ*fpn*-colonized mice exhibited a reduced ACF burden comparable to uncolonized or Δ*bft* groups, indicating that Fpn-mediated toxin activation is critical within the host tissue to facilitate ETBF-driven tumor initiation. To confirm that these phenotypes resulted specifically from loss of Fpn or BFT, we performed an independent complementation experiment in *Apc*^Min/+^ mice. Representative MB-stained distal colons demonstrated recovery of the ETBF-associated ACF phenotype with either complemented strain (Extended Data Fig. 6A), as confirmed by quantitative analysis of ACF burden (Extended Data Fig. 6B). Complementation also restored colon length (Extended Data Fig. 6C) and gross intestinal morphology toward the ETBF WT phenotype (Extended Data Fig. 6D).

**Fig. 3.**
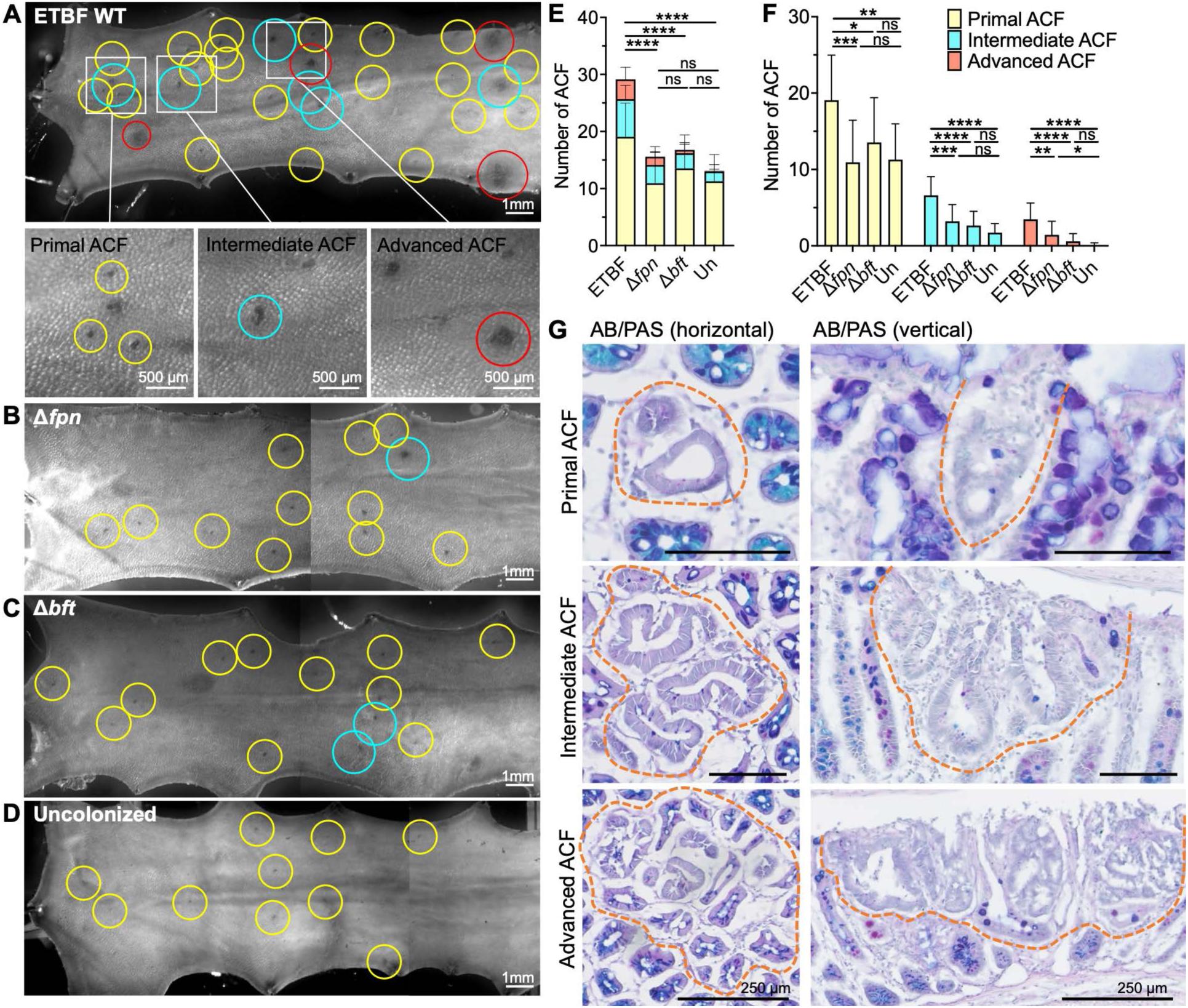
ETBF induces precancerous lesions early in life in a Fpn-and BFT-dependent manner. *Apc*^Min/+^ pups were neonatally colonized with ETBF WT, Δ*fpn*, or Δ*bft* strains or uncolonized (Un), and colons were collected at 4 weeks of age. Tissues were opened longitudinally, stained with methylene blue (MB), and imaged to visualize aberrant crypt foci (ACF). Representative MB-stained colons are shown for (A) WT, (B) Δ*fpn*, (C) Δ*bft*, and (D) uncolonized controls. In (A), boxed regions are shown below at higher magnification to illustrate representative primal (yellow circles), intermediate (blue circles), and advanced (red circles) ACF. (E) Total ACF counts and (F) distribution of ACF subtypes are quantified. (G) AB/PAS staining of representative primal, intermediate, and advanced ACF lesions in horizontal (left) and vertical (right) sections to assess mucin-depleted foci (MDF). Dashed lines outline ACF/MDF lesions. Scale bars, 100 μm unless otherwise indicated. Two to three independent biological replicates were performed; n = 15 (ETBF WT), n = 14 (Δ*fpn*), n = 17 (Δ*bft*), and n = 11 (Uncolonized). *p < 0.05, **p < 0.01, ***p < 0.001, ****p < 0.0001, ns = not significant.

The total number of ACF observed in *Apc*^Min/+^ pups was similar between 4 and 8 weeks old after neonatal ETBF colonization (Extended Data Fig. 7A), indicating that lesion initiation occurs during the early colonization window. Representative MB-stained colons (Extended Data Fig. 7B) and gross colon anatomy (Extended Data Fig. 7C) illustrate that ETBF colonization leads to increased ACF formation and architectural distortion compared with uncolonized *Apc*^Min/+^ mice by 8 weeks of age. Because progression and enlargement of individual lesions could mask changes in total ACF number, we further quantified lesions according to stage and size. To better resolve progression of larger lesions, advanced ACF were subdivided into advanced ACF (<1 mm) and macroadenomas (≥1 mm). Although total ACF burden remained similar between 4 and 8 weeks, lesion subclassification revealed a shift toward larger lesions at 8 weeks, characterized by a trend toward fewer primal ACF and a significant increase in macroadenomas (≥1 mm) (Extended Data Fig. 7D). These findings indicate that ETBF-induced lesions present at 4 weeks continue to progress in size and stage over time. To further characterize lesion morphology, AB/PAS staining of horizontal and vertical sections of distal colon tissue revealed that ETBF-induced ACF lesions exhibit features of mucin-depleted foci (MDF) (Fig. 3G, Supplementary Fig. 2), a hallmark of early epithelial transformation. In addition, epithelial cells within ACF lesions displayed elongated and densely stained nuclei compared to adjacent normal crypts, consistent with increased nuclear density (Fig. 3G). Consistent with the AB/PAS findings, Ulex europaeus agglutinin (UEA1) staining showed reduced goblet-cell associated mucin signal within ACF lesions (Supplementary Fig. 3), supporting impaired mucin production. These findings indicate that early-life ETBF colonization promotes the formation of precancerous lesions with features of early epithelial transformation.

### Regional Wnt amplification explains distal colon susceptibility to ETBF-driven neoplasia

Although ETBF has been reported to preferentially colonize the cecum in mice ^30^, ETBF-induced tumorigenic activity is most pronounced in the distal colon. Quantification of tissue-associated ETBF following extensive washing of colonic tissue specimens revealed comparable colonization across the cecum, proximal colon, and distal colon, with lower levels in the ileum (Supplementary Fig. 1), indicating that apparent cecal enrichment largely reflects luminal or mucus-associated bacteria. We therefore examined whether regional differences in epithelial physiology contribute to this bias. Wnt signaling was investigated as a candidate pathway of interest given its well-established role in crypt stem cell proliferation and adenoma initiation in colorectal cancer ^41,42,51^. Wnt reporter mice were colonized via vertical transmission with ETBF WT, Δ*fpn, or* Δ*bft* strains, then proximal colon and distal colon were evaluated for ETBF-induced Wnt activation. Reporter activity was significantly increased in the proximal (Extended Data Fig. 8A) and distal (Extended Data Fig. 8B) colon tissues of ETBF WT-colonized mice relative to that observed in the cecum (Fig. 2B). Wnt activation in the cecum and proximal colon was confined to the crypt base, however the distal colon exhibited Wnt activation extending along the crypt LP axis and also within the stromal cells underlying the colonic epithelium (Extended Data Fig. 8B). Consistent with this observation of Wnt hyperactivation, the Ki67⁺ proliferative zone in the distal colonic epithelium was expanded upon ETBF colonization (Extended Data Fig. 8C). This upward extension of the proliferative compartment has been associated with beta-catenin/Wnt-dependent tumorigenesis ^52^. As stroma-derived Wnt ligands are primary regulators of epithelial proliferation in the colon and imbalanced Wnt/APC signaling leads to upward expansion of the proliferative (stem cell) zone, promoting colon tumorigenesis ^42,52^, these results offer a potential mechanistic explanation for the pronounced neoplastic response in the distal colon. In this context, even limited ETBF colonization may be sufficient to promote substantial epithelial remodeling by amplifying stromal signaling. Thus, despite comparable tissue-associated ETBF across intestinal regions, the distal colon appears intrinsically more susceptible to Wnt-driven neoplastic transformation, explaining the regional bias in early neoplastic lesion distribution.

To define how Wnt signaling is engaged during early lesion formation, we generated Wnt reporter (TCF/Lef:H2B-GFP) × *Apc*^Min/+^ mice and performed confocal imaging of distal colon tissue at 4 weeks following neonatal ETBF colonization (Fig. 4). Even at the earliest (primal) stage, ACF lesions exhibited strong epithelial Wnt activation (Fig. 4A). Ki67 staining revealed expanded proliferative compartments extending along the crypt axis and within ACF lesions (Fig. 4B), indicating that these programs are engaged early during lesion initiation. As lesions progressed, Wnt activity showed increased spatial overlap with Ki67-positive cells (Fig. 4B). We also colonized mice at the juvenile stage (P28) and analyzed distal colon tissue at 2 months of age to assess the role of developmental timing. Under these conditions, Wnt-activated ACF lesions were not observed (Fig. 4C). Together, these results indicate that ETBF-driven tumor initiation in the distal colon is associated with early and spatially expanded epithelial Wnt activation, distinguishing this process from spontaneous tumorigenesis in *Apc*^Min/+^ mice.

**Fig. 4.**
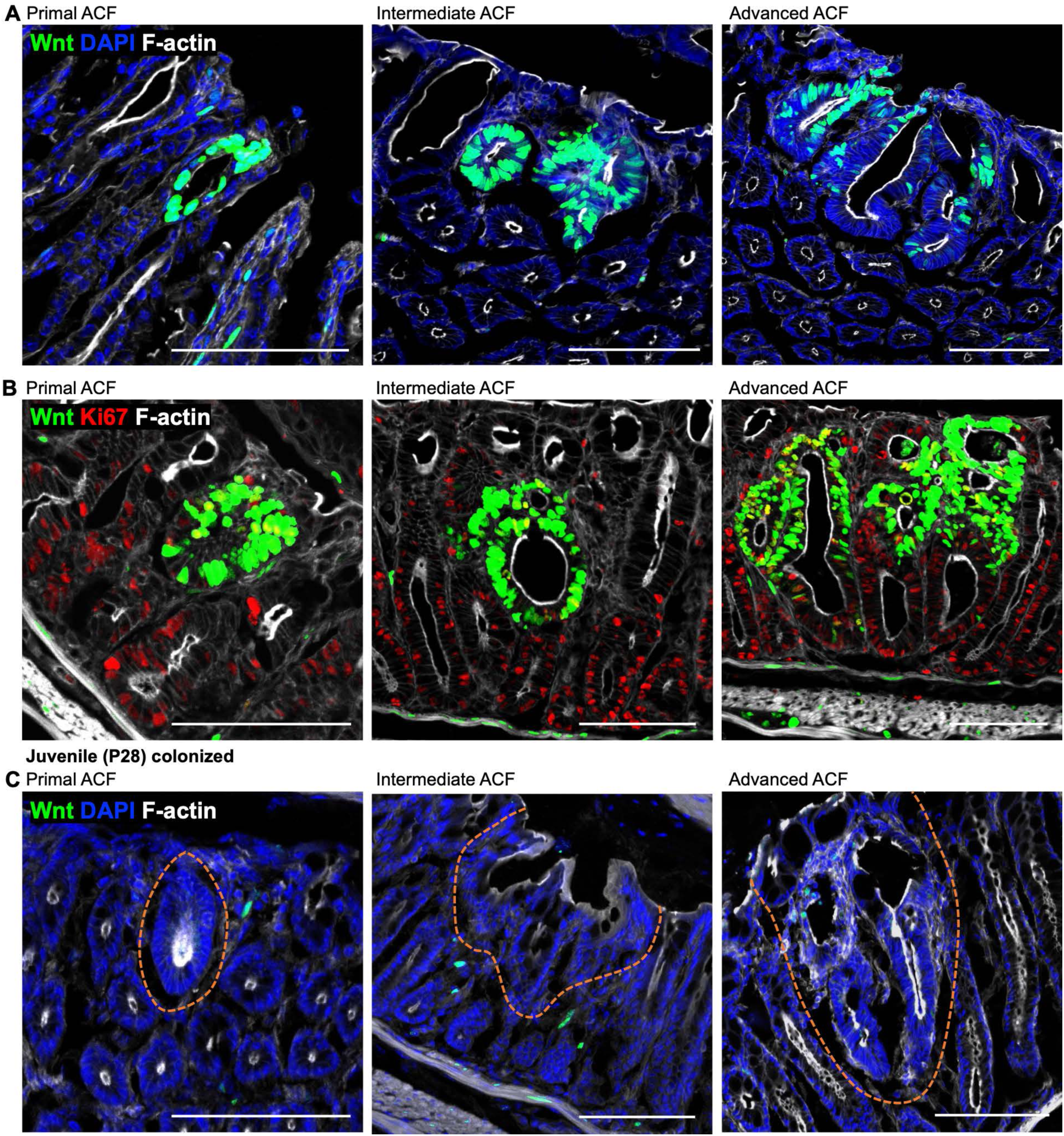
Early activation of Wnt signaling and proliferation during ETBF-induced ACF initiation. Distal colon tissue was collected 4 weeks after neonatal colonization of TCF/Lef:H2B-GFP; *Apc*^Min/+^ mice, processed using the Swiss-roll technique, and prepared for confocal imaging. (A) Representative confocal images of primal, intermediate, and advanced ACF lesions. Wnt activity (green), DAPI (blue), and F-actin (white) are shown. (B) Corresponding sections stained for Ki67 (red) together with Wnt activity (green) and F-actin (white). Increased overlap between Wnt activity and Ki67 is observed in more advanced lesions. (C) Representative confocal images from mice colonized at the juvenile stage (P28) and analyzed at 2 months of age show absence of Wnt^+^ ACF lesions. Scale bars, 100 μm. Scale bars, 100 μm.

### Early-life ETBF colonization drives intracellular bacterial community formation within tumor lesions

Human studies have shown that ETBF frequently resides within invasive biofilms on the preneoplastic colonic mucosa of patients with familial adenomatous polyposis (FAP), highlighting a strong association with tumor initiation and promotion ^26^. Building on these observations, we asked whether early-life ETBF colonization in our *Apc*^Min/+^ model might similarly permit ETBF to co-localize within ACF lesions. To enhance analysis of ETBF co-localization in ACF lesions, 8-week-old mice were used, in which lesions were larger and more readily identifiable. Using the Swiss-roll technique ^53^ to survey the distal colon, we found that ETBF formed discrete intracellular bacterial communities (IBCs) within tumor lesions (Fig. 5A). Orthogonal views and video analysis of z-stack confocal images confirmed that these intramural bacterial aggregates represent *de novo* intracellular colonization (Extended Data Fig. 9, Movie 1-2). Δ*fpn* and Δ*bft* strains were likewise localized exclusively to tumor lesions, IBC formation was reduced in both mutants, with Δ*fpn* displaying an intermediate phenotype and Δ*bft* showing a more pronounced reduction relative to ETBF WT (Fig. 5B-C). This graded reduction in IBC formation parallels the intermediate tissue colonization and remodeling phenotypes observed with Δ*fpn* and the stronger attenuation seen with Δ*bft* earlier in the study (Fig. 1B-E), supporting the idea that spatial activation of BFT facilitates epithelial invasion and intramural expansion.

**Fig. 5.**
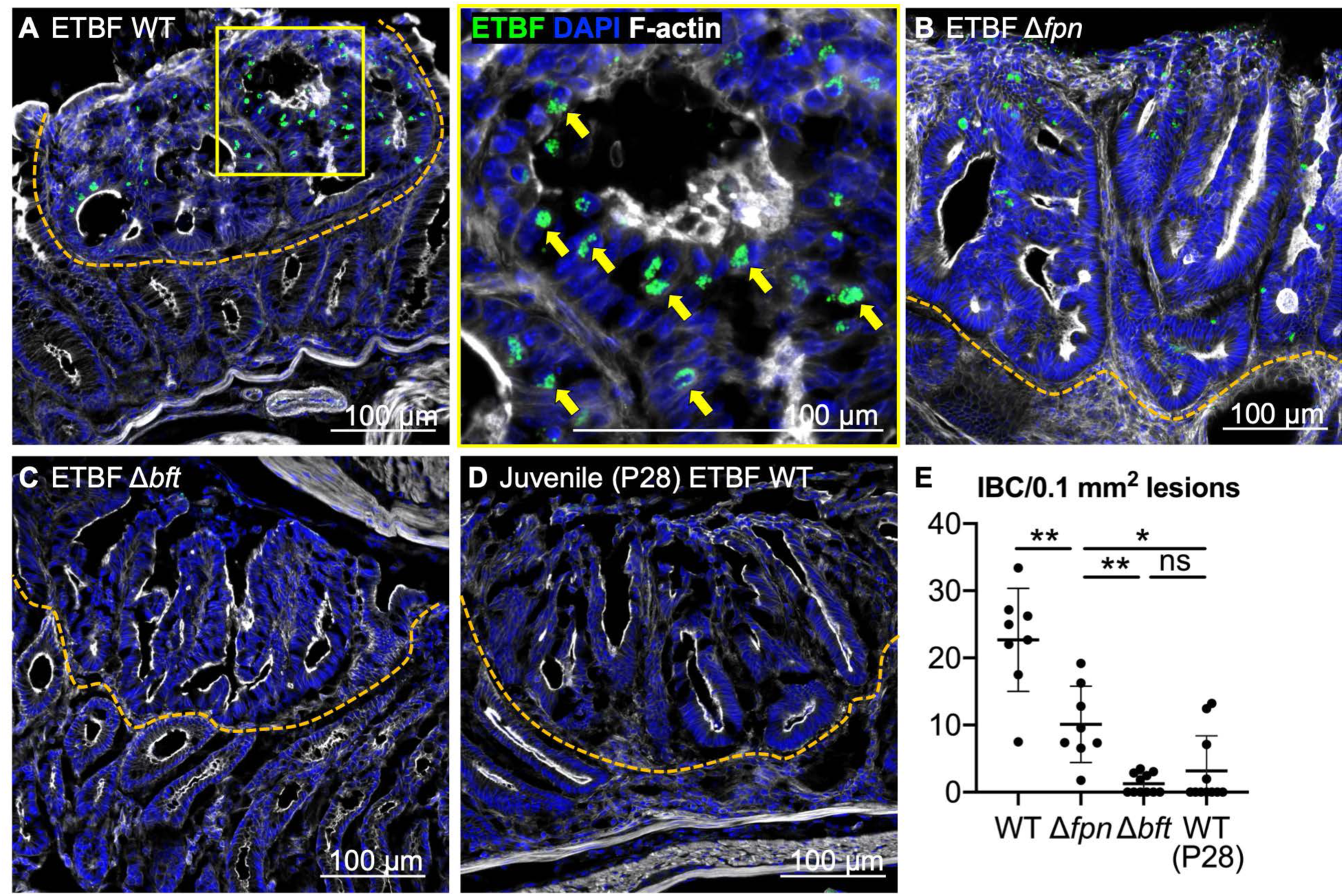
ETBF establishes toxin-and developmental timing-dependent intracellular bacterial communities (IBCs) within early tumor lesions. Confocal imaging of distal colons from 8-week-old *Apc*^Min/+^ mice neonatally colonized with (A) ETBF WT, (B) Δ*fpn*, or (C) Δ*bft*, and (D) *Apc*^Min/+^ mice colonized with ETBF WT at the juvenile stage (P28). ETBF (green), DAPI (blue), and F-actin (white) are shown. Dashed lines outline ACF lesions. The yellow box in (A) denotes the region shown at higher magnification; yellow arrows indicate IBCs. (E) Quantification of IBCs per 0.1 mm² of lesion area. Scale bars, 100 μm.

To assess whether IBC formation was dependent on the host developmental stage at which ETBF was introduced into the colonic microbiome, colonization was initiated in mice at P28 (juvenile stage). IBCs were markedly reduced following juvenile colonization (Fig. 5D). Quantification of IBCs normalized to lesion area confirmed these patterns, with Δ*fpn* showing an intermediate reduction, whereas both Δ*bft* and juvenile ETBF WT showed markedly lower IBC levels relative to neonatally colonized ETBF WT (Fig. 5E). Thus, later colonization substantially limits IBC formation, indicating that early-life exposure strongly favors the establishment of these intracellular communities. This finding also argues that ongoing luminal exposure alone is insufficient to account for the robust IBC formation observed following neonatal colonization and is consistent with a contribution from lamina propria niche acquisition, which is temporally restricted to the pre-weaning period ^30^. Together, these findings suggest that ETBF persistence within transformed epithelium may reinforce oncogenic signaling, positioning ETBF not only as an initiator of precancerous change in early life but also as a microbe capable of adaptation to the tumor microenvironment.

### Developmental window NTBF colonization prevents ETBF-driven tumor initiation

We previously showed that colonization with non-toxigenic *B. fragilis* (NTBF) during the postnatal day 12-16 (P12-16) developmental window effectively prevents ETBF from accessing the colonic LP niche ^30^. To test whether exclusion of ETBF from the LP alters tumor initiation, we pre-colonized *Apc*^Min/+^ pups with NTBF. Colonization at P12, but not P21, significantly reduced ETBF-induced ACF formation (Fig. 6A-B). This protection required the NTBF Type VI Secretion System (T6SS), as a Δ*tssC* strain failed to block ACF formation (Fig. 6C), consistent with prior evidence that T6SS-dependent competition underlies NTBF exclusion of ETBF from the tissue niche ^30^. Quantification of ACF burden across conditions (Fig. 6D-E) demonstrated that early NTBF colonization at P12 provides significantly greater protection compared to P21 intervention. In addition, the Δ*tssC* mutant failed to confer protection, with ACF levels comparable to ETBF only controls, indicating that T6SS-mediated interbacterial competition is essential for effective exclusion of ETBF and prevention of ETBF-driven tumor initiation. Confocal imaging confirmed that NTBF-mCherry restricted ETBF-sfGFP localization to the mucus layer in P12-protected mice, whereas ETBF-sfGFP predominated and showed deeper penetration in P21-protected animals (yellow arrow, Extended Data Fig. 10A). Importantly, ETBF established IBCs within ACF lesions in the P21-protected group, whereas no IBCs were found in P12-protected group, indicating successful protection (Fig. 6F). Bacterial burden analysis further showed that NTBF pre-colonization at P12 reduced ETBF colonization in the lumen compared to P21 NTBF and P12 NTBF Δ*tssC* (Extended Data Fig. 10B). Pathology analysis supported these findings, as P12 NTBF colonization improved colon length (Extended Data Fig. 10C), cecal weight (Extended Data Fig. 10D), and colon appearance (Extended Data Fig. 10E), whereas P21 NTBF or P12 NTBF *ΔtssC* conferred no protection. These findings extend prior work on developmental window colonization dynamics by showing that NTBF not only excludes ETBF from its pathogenic niche but also prevents its carcinogenic sequelae *in vivo*.

**Fig. 6.**
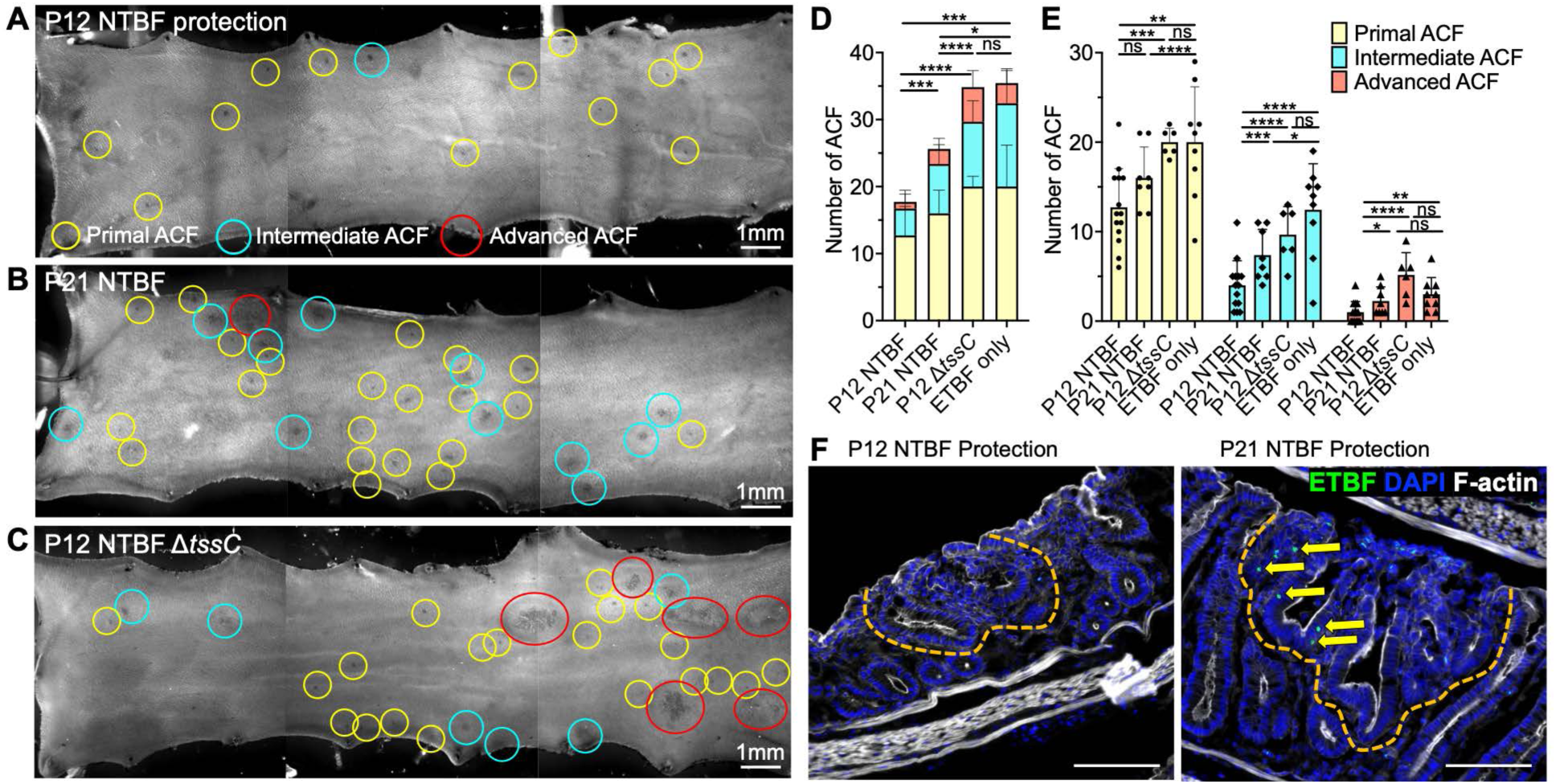
Early-life NTBF colonization protects against ETBF-driven tumor initiation in a T6SS-dependent manner. (A-C) Representative MB-stained colons from *Apc*^Min/+^ pups born to dams colonized with ETBF WT and gavaged with (A) NTBF at postnatal day 12 (P12), (B) NTBF at P21, or (C) NTBF Δ*tssC* at P12. ACF were classified as primal (yellow circles), intermediate (blue circles), or advanced (red circles). (D) Quantification of total ACF across all groups including P12 NTBF, P21 NTBF, P12 Δ*tssC*, and ETBF only controls. (E) Distribution of ACF subtypes (primal, intermediate, and advanced) across all groups is shown. (F) Confocal imaging of ACF lesions in P12-and P21-protected mice. Intracellular bacterial communities (IBCs; yellow arrows) are observed in P21-protected mice but are absent in P12-protected mice. Dashed lines outline ACF lesions. Scale bars, 100 μm unless otherwise indicated. Two independent replicates were performed; n = 14 (P12 NTBF), n = 8 (P21 NTBF), n = 6 (P12 Δ*tssC*), and n = 11 (ETBF only). *p < 0.05, **p < 0.01, ***p < 0.001, ****p < 0.0001, ns = not significant.

## Discussion

Our findings extend the paradigm of microbe-cancer interactions by identifying developmental timing as a key determinant of colorectal cancer (CRC) susceptibility. As early-life microbial exposures are increasingly recognized as critical determinants of lifelong health, this study provides one of the first mechanistic demonstrations of how a pioneer microbial colonizer can durably remodel epithelial cell states to predispose the colon to cancer. Using a vertical transmission model that preserves the native microbiota, we show that early-life acquisition of the lamina propria niche by enterotoxigenic *Bacteroides fragilis* (ETBF) drives precancerous changes in the colon. This process requires both expression of the metalloprotease toxin BFT and its activation by Fragipain (Fpn), underscoring that localized toxin activity within the tissue niche is critical to induce the epithelial stem cell remodeling and early tumor initiation. This compartment-specific requirement is consistent with prior work showing that colonic mucus-associated protease activity can mediate BFT cleavage in the intestinal lumen, whereas Fpn is required for BFT activation in non-luminal settings such as bloodstream infection. We therefore propose that after early-life ETBF gains access to the lamina propria, where exposure to luminal mucus-associated protease activity is limited, Fpn-dependent local toxin activation becomes important for sustained epithelial remodeling.

Building on prior models of ETBF-induced tumorigenesis, which have provided key insights into colonization dynamics in juvenile or adult *Apc*^Min/+^ mice ^54,55^, our study identifies an additional developmental phase of host-microbe interaction that has not been previously examined. The vertical transmission model captures an early window when the lamina propria is permissive to microbial entry and epithelial fate programs remain plastic, allowing ETBF to reside in close proximity to crypt stem cells and induce localized toxin-dependent remodeling. Although early-life colonization by ETBF has generally been considered asymptomatic, our findings reveal that neonatal exposure incites durable remodeling of colonic epithelium, establishing a long-term vulnerability that increases colorectal cancer susceptibility. Notably, ETBF colonization initiated in juvenile mice did not produce detectable tumor-associated IBCs, whereas neonatal colonization resulted in IBC-positive ACF lesions. This timing-dependent difference suggests that early-life ETBF colonization creates a permissive antecedent state for later tumor-associated intracellular localization. One possibility is that early establishment of a lamina propria tissue niche increases the likelihood of subsequent ETBF persistence within the tumor microenvironment. These findings suggest that developmental timing shapes ETBF pathogenesis and its interaction with the host epithelium within the tumor microenvironment. However, whether tumor-associated IBC formation directly contributes to epithelial reprogramming or tumor progression remains to be determined. Although these findings are consistent with a developmental window of susceptibility, our neonatal versus juvenile colonization experiments primarily test whether ETBF gains access to the early-life LP niche, rather than directly testing whether neonatal epithelial cells are intrinsically more susceptible than juvenile or adult epithelial cells to Wnt activation. Distinguishing epithelial cell-intrinsic age-dependent susceptibility from the effects of bacterial tissue localization will require future approaches that uncouple host epithelial developmental stage from ETBF access to the lamina propria niche.

A central insight of this work is that ETBF-induced oncogenic responses are regionally distinct. Although tissue-associated ETBF colonization is comparable across the cecum, proximal colon, and distal colon, tumor precursors predominantly arise in the distal colon. This regional uncoupling highlights the role of tissue context in dictating susceptibility to ETBF-induced ACF initiation. We show that early-life ETBF colonization amplifies Wnt hyperactivation in the distal colon, providing a mechanistic explanation for why this region is uniquely prone to neoplastic transformation. Notably, in a susceptible mouse model, early ACF lesions exhibited strong epithelial Wnt activity in the distal colon when neonatally colonized with ETBF, supporting a role for ETBF in promoting Wnt-associated early tumorigenesis. In contrast, Wnt-activated ACF lesions were not observed following juvenile colonization, further supporting a role for Wnt-mediated mechanisms in ETBF-induced ACF initiation. The marked rise of early-onset CRC in the descending (left) colon and rectum in humans ^6,7^ parallels our observation that ETBF-induced precancerous lesions predominantly develop in the distal colon. This concordance suggests that early-life microbial toxin exposure may contribute to the regional bias of human disease by amplifying stromal and epithelial Wnt signaling within the distal colonic microenvironment. Given the well-established role of Wnt signaling in colorectal tumorigenesis, our findings support a model in which early-life ETBF colonization promotes Wnt-mediated pathways to drive ACF initiation. However, the dependency upon Wnt signaling, the relative contributions of stromal versus epithelial Wnt activation, and the temporal sequence linking ETBF colonization, Wnt activation, and ACF initiation remain unresolved. Future studies incorporating functional perturbation and longitudinal analyses will be required to further define the temporal sequence and mechanistic basis of these events.

Taken together, neonatal ETBF colonization thus initiates a continuum of lamina propria-associated pathogenesis that extends from early epithelial remodeling to tumor-associated persistence. We observe that ETBF persists within tumor lesions as organized intracellular communities, reminiscent of *Fusobacterium nucleatum* in human CRC ^56^ and *H. pylori* in gastric cancer ^57^. Unlike other CRC-associated bacteria such as *F. nucleatum* or *Peptostreptococcus anaerobius* ^56,58^, which are typically detected as opportunistic or strain-specific colonizers in established tumors, ETBF is a prevalent early-life colonizer of the human gut. By occupying this developmental niche, ETBF promotes early epithelial remodeling and establishes persistent intramural residency within precancerous lesions. The discovery of tumor-associated ETBF IBCs highlights important future directions, including determining whether these communities remain functionally active via sustained BFT expression, how they shape host-microbe interactions during tumor development, and whether analogous IBCs or tissue-associated microbial niches can be detected in human colorectal tissues.

These findings also align with recent genomic analyses of human colorectal cancers, which revealed that mutational signatures caused by microbiome-derived genotoxins such as colibactin are enriched in early-onset cases and are often imprinted early during tumor evolution ^59^. Our findings provide a mechanistic explanation for these observations, showing that early-life microbial toxin exposure can durably remodel epithelial cell states and establish a long-lasting oncogenic field that predisposes to malignant transformation. However, as these studies were performed in a genetically predisposed *Apc*^Min/+^ model, it remains to be determined how ETBF-induced epithelial remodeling influences tumor initiation and progression in non-FAP contexts. Addressing this will require models that couple early-life microbial exposure with subsequent tumor initiation. At the same time, this model provides a tractable system to interrogate early tumor-initiating events, enabling resolution of how early-life microbial exposure shapes epithelial susceptibility prior to overt malignancy. This work positions ETBF as a model for how developmental microbial exposure can influence both the initiation and progression of colorectal cancer.

Finally, our observation that non-toxigenic *B. fragilis* (NTBF) competitively restricts ETBF colonization and ETBF-induced early tumorigenesis during this developmental window underscores how commensal community structure established during a critical early-life window can shape cancer susceptibility. These results point to the possibility that targeted modulation of early-life microbiota could serve as a preventive strategy against toxin-mediated microbial carcinogenesis.

## Methods

### Bacterial strains, culture conditions, and antibiotics

The bacterial strains used in this study are listed in Supplementary Table 1. *Bacteroides fragilis* strains were routinely cultured anaerobically in brain heart infusion medium supplemented with hemin (0.0005%) and vitamin K₁ (0.5 µg/ml) at 37 °C in a Coy anaerobic chamber with a gas mixture of 5% H₂, 10% CO₂, and 85% N₂. *Escherichia coli* S17-1 λpir was used for plasmid construction and conjugation into *B. fragilis*. Fluorescent and mutant *B. fragilis* derivatives (Supplementary Table 1) were generated and sequence-verified following established procedures (*31*). All enterotoxigenic *B. fragilis* (ETBF) strains were engineered to constitutively express superfolder GFP (sfGFP) from a pNBU2 integrative vector under control of a phage promoter, and non-toxigenic *B. fragilis* (NTBF) strains were engineered to express mCherry from the analogous vector (*30*). When necessary, antibiotics were used at standard working concentrations: 100 µg ml⁻¹ ampicillin, 200 µg ml⁻¹ gentamicin, 5 µg ml⁻¹ clindamycin, 10 µg ml⁻¹ tetracycline, and 20 µg ml⁻¹ rifampicin.

### Primary colonic epithelial cell isolation and culture

Cecal tissue from 8-week-old C57Bl/6J mice was isolated, minced, and incubated in collagenase type I at 37 °C with gentle rocking and intermittent mechanical dissociation. The resulting cell suspension was filtered through a 100 μm strainer, washed, and epithelial units were collected. Cells were embedded in Matrigel (BD Biosciences) and cultured in 50% L-WRN conditioned medium diluted in primary culture medium supplemented with 10 mM Y-27632 and 10 mM SB431542 (R&D System) *(33)*. Medium was changed every 2 days, and organoids were passaged every 3 days (1:4-1:8 split). For toxin treatment, organoids were incubated with 300 ng/ml activated BFT or inactive BFT for the indicated time points prior to imaging.

### Animal studies

All mouse experiments were conducted in accordance with ethical regulations and approved protocols of the Institutional Animal Care and Use Committee (IACUC) and Institutional Biosafety Committee (IBC) at Washington University School of Medicine. C57Bl/6J mice and all genetically modified lines used in this study were obtained from The Jackson Laboratory and bred in-house (Supplementary Table 2). Mice were maintained under specific-pathogen-free (SPF) conditions (20-26 °C, 30-70% humidity, 12 h light/dark cycle) with ad libitum access to standard chow and autoclaved water. Sample size estimates for all experiments were based on prior colonization studies performed within the laboratory (*31*). Breeding pairs were established to generate experimental cohorts, and both male and female pups were used.

### Neonatal colonization model

Vertical transmission experiments were used to model early-life colonization by *B. fragilis*. Briefly,

*B. fragilis* strains were cultured anaerobically overnight to stationary phase (16∼18 h) in 35 ml of brain heart infusion medium supplemented with hemin and vitamin K₁ (BHIS), as described above. Cells were pelleted by centrifugation at 5,000 × g for 10 min at 4 °C, washed, and resuspended in 0.1 N sodium bicarbonate. Pregnant dams (8-10 weeks old) received 100 mg l⁻¹ clindamycin in drinking water from embryonic day 14 (E14) to E17. On E15, dams were gavaged with 100 µl of the prepared inoculum containing 5 × 10⁹ CFU of *B. fragilis* to establish maternal colonization. Offspring were delivered naturally and remained with the colonized dam. Stool samples from dams and pups were collected at multiple time points to monitor colonization stability and density. This model recapitulates vertical transmission and persistent colonization during the neonatal window, enabling investigation of early-life microbial programming (*30*). For all *in vivo* experiments, no investigator blinding was applied during animal handling or outcome assessment. Animals were excluded only if the corresponding dam was insufficiently colonized (<10⁸ CFU g⁻¹ of *B. fragilis* in fecal samples) to support vertical transmission. Mice were euthanized by CO₂ inhalation followed by cervical dislocation in accordance with approved institutional animal welfare protocols.

### *In vivo* probiotic competition studies

To evaluate the ability of probiotic NTBF to competitively inhibits ETBF tissue colonization during early-life and prevent ETBF-induced precancerous lesion development, neonatal mice colonized vertically with ETBF were subjected to postnatal probiotic interventions. Pups were gavaged with 50 µl of 0.1 N sodium bicarbonate containing 5 × 10⁹ CFU of either NTBF or its isogenic Δ*tssC* mutant on postnatal day (P) 12, or P21. All animals were euthanized on P28 for tissue collection and quantitative analyses of bacterial colonization and precancerous lesion formation.

### Juvenile colonization model

To model *B. fragilis* colonization after GAPs closure, 4-week-old mice were treated with 100 mg l⁻¹ clindamycin in drinking water for 24 h to transiently reduce commensal flora. Mice were then gavaged with 100 µl of 0.1 N sodium bicarbonate containing 5 × 10⁹ CFU of either ETBF or NTBF. Following gavage, mice continued to receive 100 mg l⁻¹ clindamycin for an additional 48 h, after which antibiotic water was replaced with sterile water. Animals were euthanized 4 weeks post-colonization for endpoint analyses including CFU enumeration, histopathology, and precancerous lesion assessment.

### Tissue collection and bacterial quantification

Mice were euthanized between postnatal day (P) 21 and P60, depending on the experimental design. Colons were collected, photographed for gross pathology and length measurements, and processed for bacterial quantification and histological analyses. For niche-specific CFU analysis, ceca were opened, rinsed with PBS, and mucus was scraped from the epithelial surface. Cecal tissue, mucus, and feces were separately placed into pre-weighed sterile 2 ml screw-cap tubes containing 1 ml of sterile PBS and 1 mm or 2.3 mm beads. Homogenization was performed twice in an MP Biomedicals FastPrep 24 at 6 m s^-1^ for 60 s at 4 °C, with a 5 min cool down between homogenizations. Homogenates were serially diluted and plated for CFU enumeration on BHIS agar containing gentamicin and clindamycin, with tetracycline or rifampicin added for differential selection when appropriate. CFU counts were log_10_-transformed before plotting with a detection limit of 10³ CFU g⁻¹.

### Histopathology staining and imaging

Cecum, proximal, and distal colon tissues were fixed in 4% paraformaldehyde (PFA) in PBS, embedded in optimal cutting temperature (OCT) compound (Sakura Finetek), and cryosectioned for Alcian blue/Periodic acid-Schiff (AB/PAS) staining. AB/PAS staining was performed using the 91022B AB/PAS Stain Kit (Newcomer Supply). Whole-slide scans of AB/PAS-stained sections were generated using a Hamamatsu NanoZoomer S360 digital slide scanner, and histopathological features were analyzed with NDP.view2 software to quantify goblet cells, crypt lengths, and mucin-depleted foci (MDF).

### Immunofluorescent staining

Whole colons were fixed in 4% paraformaldehyde (PFA) in PBS for 48 h at 4 °C, cryoprotected in 20% sucrose for 48 h, frozen in liquid nitrogen, and embedded in OCT compound. Blocks were stored at −80 °C until sectioning. Sections (7 or 30 μm) were mounted on Superfrost Plus slides and stained with antibodies and probes listed in Supplementary Table 3. Samples were counterstained with ProLong Glass Antifade containing NucBlue (DAPI) and Alexa Fluor 555-or 647-phalloidin, and cured at room temperature in the dark for 72 h before imaging.

### Automated image analysis and cell quantification

Quantification of Wnt-active and Atoh1^+^ cells was performed using an automated Fiji-based pipeline. Because these signals are nuclear-localized, automated detection was used to identify positive cells. Epithelial crypt regions were defined as regions of interest, and positive cells were identified using consistent fluorescence intensity thresholding across all samples. Particle analysis (size 10-100 µm², circularity >0.1) was applied to exclude debris, and identical parameters were used across all conditions. Multiple fields from independent animals were analyzed.

### Swiss-roll preparation and cryosectioning

To visualize precancerous lesions and epithelial architecture across the entire colon, tissues were processed using a modified Swiss-roll technique (52). Immediately after dissection, colons were opened longitudinally along the mesenteric border, rinsed gently in cold PBS, and flattened mucosal side up on a moist surface. Starting from the distal end, the tissue was rolled toward the proximal end with the mucosal surface facing inward, embedded in OCT, rapidly frozen on dry ice, and stored at −80 °C. Frozen blocks were cryosectioned at 7 or 30 μm and mounted on Superfrost Plus slides for AB/PAS staining or immunofluorescence. This configuration allowed comprehensive visualization of colonic length and spatial distribution of early precancerous lesions within a single section.

### Confocal microscopy and image processing

Immunofluorescently labeled tissues were imaged using a Zeiss LSM 880 Airyscan confocal microscope equipped with 20× and 40× Plan-Apochromat objectives. Images were acquired in sequential scanning mode using appropriate excitation and emission filters for NucBlue (DAPI), sfGFP, mCherry, Alexa Fluor 555, and Alexa Fluor 647. Z-stack images were collected at 0.3 μm intervals and reconstructed into three-dimensional renderings using Zeiss ZEN Black and Volocity software. Image processing, including background subtraction, Gaussian filtering, and intensity normalization, was performed using ZEN Blue and ImageJ. Identical acquisition and processing settings were applied across all conditions. Three-dimensional movies (Supplementary Movies 1-2) were rendered using Volocity software.

### Methylene blue staining and aberrant crypt foci (ACF) counting

To assess precancerous lesion formation after neonatal colonization, *Apc*^Min/+^ mice were infected with wild-type ETBF, Δ*fpn*, or Δ*bft* mutant strains, and colons were collected at 4 weeks of age. The distal half of each colon was excised, flushed gently with cold phosphate-buffered saline (PBS), opened longitudinally along the mesenteric border, and pinned mucosal side up on SYLGARD™ 184 silicone gel. Samples were fixed flat in 4% paraformaldehyde (PFA) for 24 h at 4 °C, rinsed in PBS, and stained with 0.05% methylene blue for 5-10 min, followed by brief PBS washing to remove excess dye. Stained colons were imaged under a dissecting microscope using transmitted-light illumination. Aberrant crypt foci (ACF) were identified based on established morphological criteria, including enlarged or distorted crypts, thickened epithelial borders, and enhanced pericryptal methylene blue uptake. ACFs were classified as primal, intermediate, or advanced lesions according to crypt size, shape distortion, and multiplicity. The number and subtype of ACFs were quantified for each sample.

### Experimental design and statistical analysis

Sample sizes were estimated based on prior studies and power calculations (α = 0.05, power = 0.9), indicating that approximately 5-8 mice per group would be sufficient. To account for biological variability, a target of 8-10 mice per group was used when feasible; final group sizes varied depending on breeding yield and genotype availability. Exact sample sizes and replicate numbers are reported in the corresponding figure legends. Both male and female offspring were included, and littermate-matched controls were used in all comparisons. Data were analyzed using GraphPad Prism 10. Unpaired two-tailed Student’s t tests (or Welch’s correction when variances were unequal) and one-way ANOVA with Tukey’s or Dunnett’s multiple-comparison tests were used as appropriate. Error bars represent standard deviation (SD), and P < 0.05 was considered statistically significant.

## Acknowledgments

We thank members of the Bubeck-Wardenburg laboratory for helpful discussions and technical assistance. This work was supported by the National Institutes of Health Pediatric Gastroenterology Research Training Program (T32 DK077653 to S.K.R. and J.J.P.), a National Institutes of Health/National Cancer Institute postdoctoral fellowship (F32CA306217-01 to S.K.R.), the National Institutes of Health Medical Scientist Training Program at Washington University (T32 GM007200 to J.J.P.), a National Institutes of Health/National Institute of Allergy and Infectious Diseases predoctoral fellowship (F30AI197460 to J.J.P), and a National Institutes of Health research grant (R01AI138565 to J.B.W.).

## Author contributions

S.K.R. and J.B.W. conceptualized the project. S.K.R., E.V. and J.B.W. developed the methodology. S.K.R., E.V. and J.J.P. conducted the investigations. S.K.R., E.V. and J.J.P. performed visualization. S.K.R., J.J.P. and J.B.W. acquired funding. S.K.R. wrote the original draft. S.K.R., E.V., J.J.P. and J.B.W. reviewed and edited the manuscript.

## Competing interests

Authors declare that they have no competing interests.

## Data and materials availability

All data are available in the main text or the supplementary materials. Bacterial strains, plasmids, and mouse lines generated in this study are available from the corresponding author under a material transfer agreement with Washington University.

## Additional information

Supplementary information:

Supplementary Figs. 1-3

Supplementary Tables 1-3

Supplementary Movies 1-2

---

**Extended Data Fig. 1.**
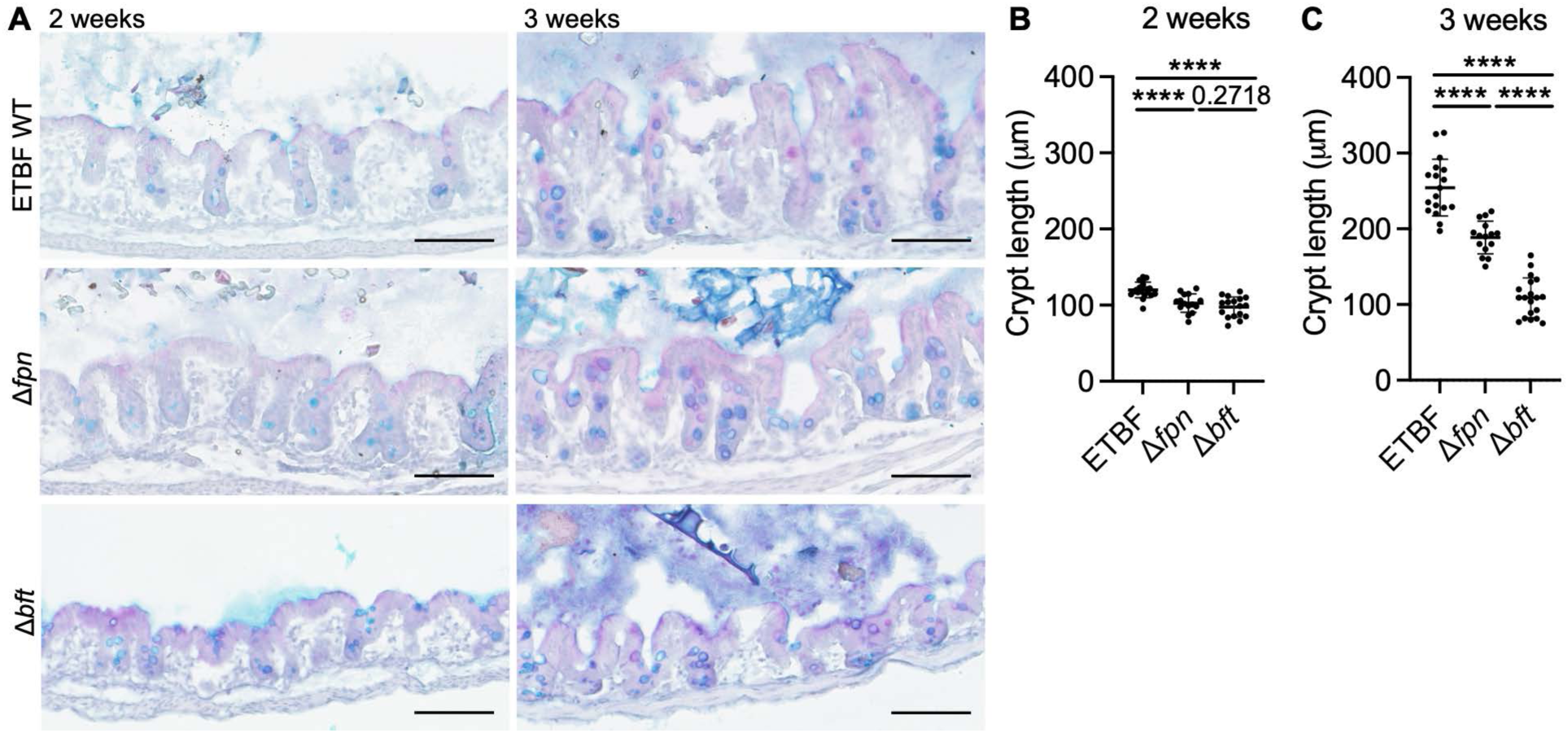
Early-life developmental dynamics of ETBF-induced crypt remodeling, related to. Fig. 1. (A) Representative AB/PAS staining of cecal tissue from mice neonatally colonized with ETBF WT, Δ*fpn*, or Δ*bft* strains at 2 and 3 weeks of age. (B-C) Quantification of crypt length at (B) 2 and (C) 3 weeks of age. Data are presented as mean ± SD. ****p < 0.0001.

**Extended Data Fig. 2.**
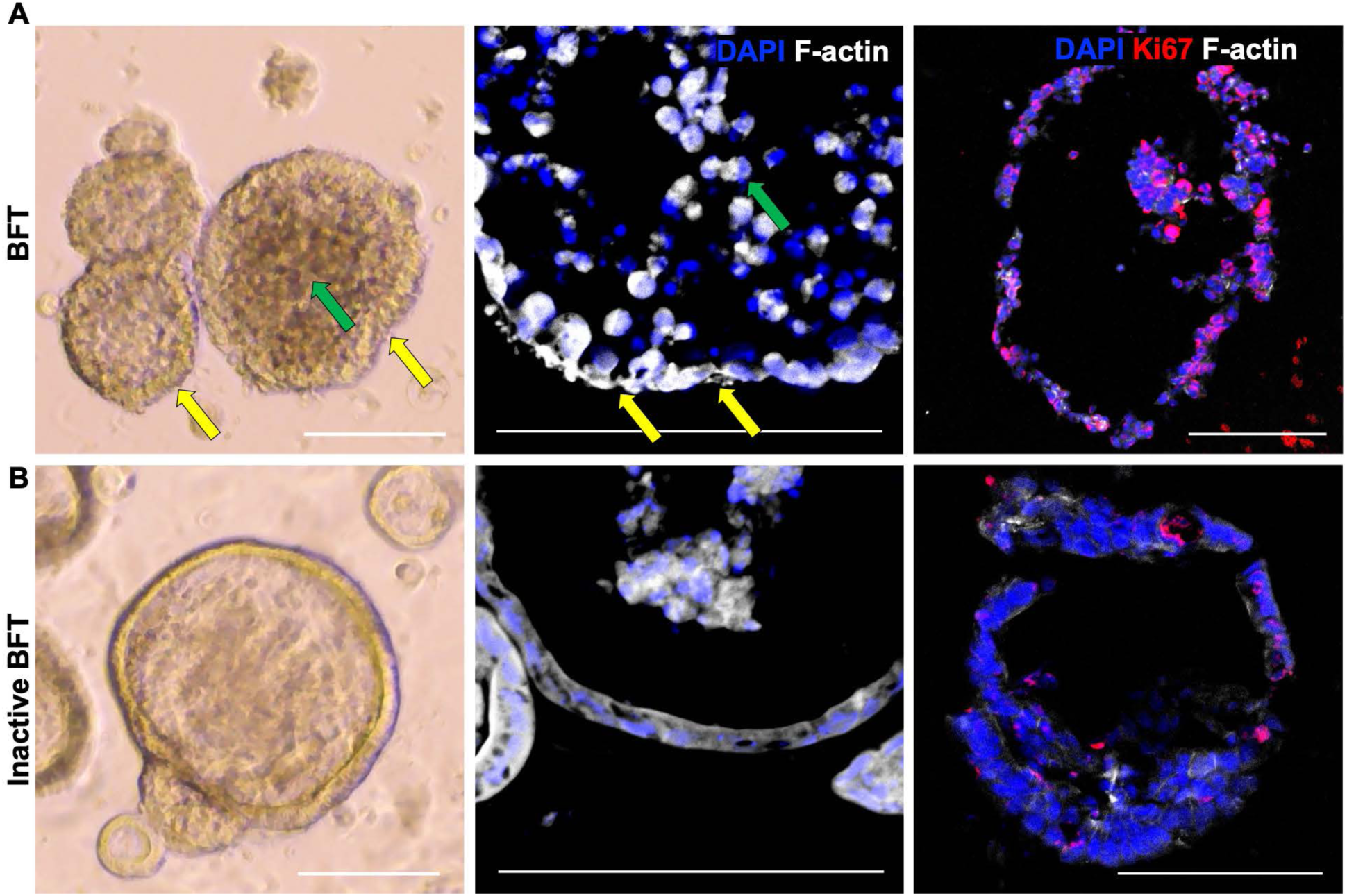
BFT induces rapid epithelial remodeling and increased proliferative activity in primary colonic organoids, related to. Fig. 1. Primary colonic epithelial organoids derived from C57Bl/6J mice were treated with 300 ng/ml of activated BFT (A) or inactive BFT (B) for 3 hours. Left, representative brightfield images; green arrows indicate dense cellular centers and yellow arrows indicate disrupted epithelial junctions. Middle, representative confocal images of DAPI (blue) and F-actin (white), highlighting epithelial disorganization and surface disruption. Right, representative confocal images of DAPI (blue), Ki67 (red), and F-actin (white). Scale bars, 100 µm. Scale bars, 100 µm.

**Extended Data Fig. 3.**
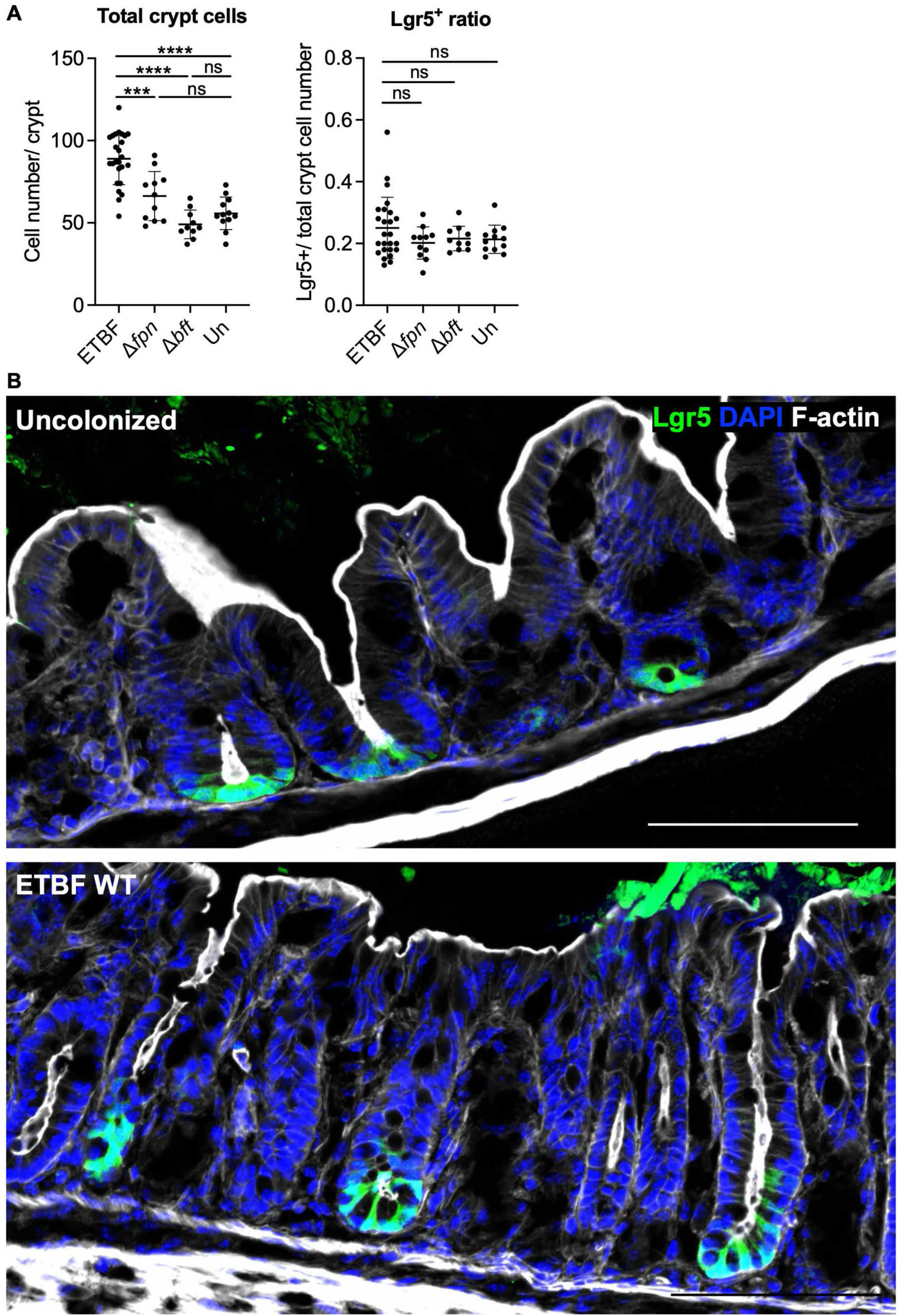
Crypt cell quantification in Lgr5 reporter mice, related to. Fig. 2. Lgr5 reporter mice were neonatally colonized with ETBF WT, Δ*fpn*, or Δ*bft* strains, or left uncolonized, and analyzed at 3 weeks of age. (A) Total cell number per crypt (left) and the ratio of Lgr5⁺ to total crypt cells (right) were quantified in the cecum. (B) Representative confocal images of uncolonized and ETBF WT-colonized cecal crypts, shown at larger scale for improved visualization of Lgr5⁺ cells. Data are shown as mean ± SD. ***p < 0.001, *\*\*\*\**p < 0.0001, ns = not significant.

**Extended Data Fig. 4.**
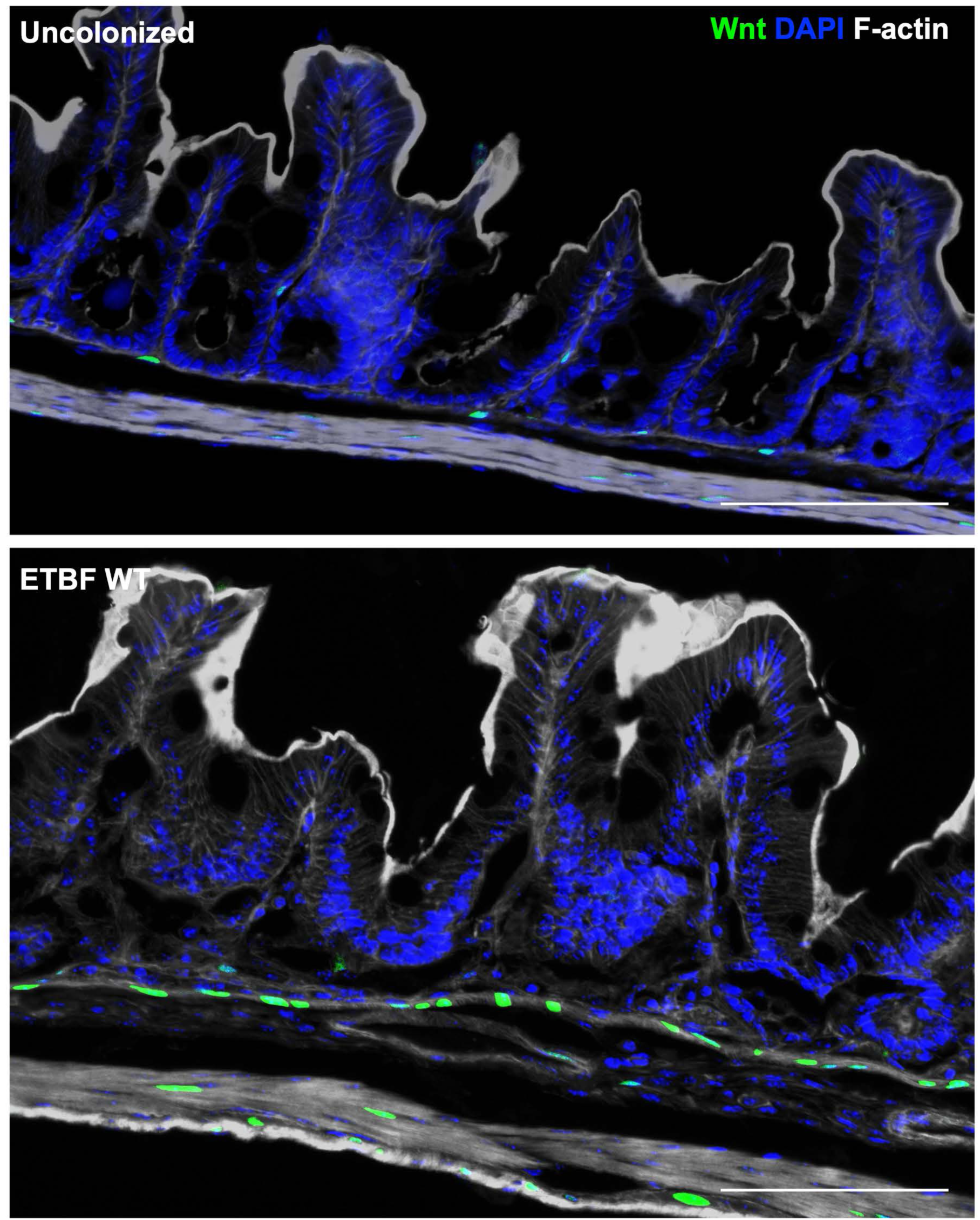
Enlarged views of Wnt activity reporter in cecal crypts, related to Fig. 2. TCF/Lef:H2B-GFP reporter mice were colonized with ETBF WT or left uncolonized and analyzed at 3 weeks of age. Representative confocal images of cecal crypts are shown at larger scale to improve visualization of Wnt activity. Scale bars, 100 μm.

**Extended Data Fig. 5.**
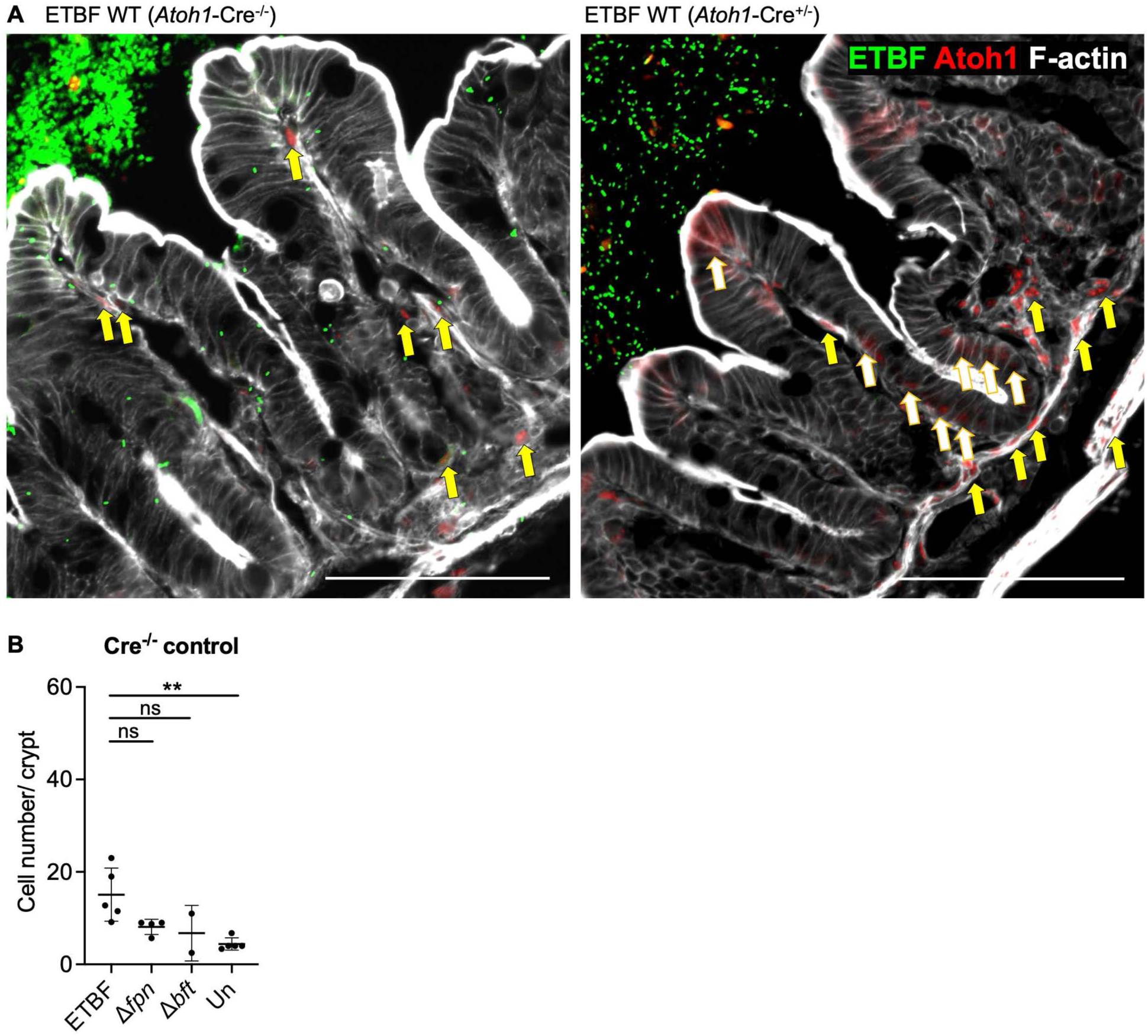
Validation of Atoh1 reporter specificity, related to. Fig. 2. R26-LSL-H2B-mCherry; Atoh1-Cre^-/-^ littermate controls were neonatally colonized with ETBF WT, Δ*fpn*, or Δ*bft* strains, or left uncolonized, and analyzed at 3 weeks of age. (A) Representative confocal images comparing ETBF WT-colonized Atoh1-Cre⁻^/^⁻ (left) and Atoh1-Cre⁺^/^⁻ (right) tissues. Stromal mCherry⁺ cells are indicated by yellow arrows in both panels, while epithelial mCherry⁺ cells are indicated by white-filled yellow arrows in the Atoh1-Cre⁺^/^⁻ panel (right). (B) Quantification of mCherry⁺ cells per cecal crypt. Differences in stromal background likely reflect expansion of the stromal compartment associated with crypt elongation, rather than reporter-specific signal. Data are shown as mean ± SD. **p < 0.01, ns = not significant.

**Extended Data Fig. 6.**
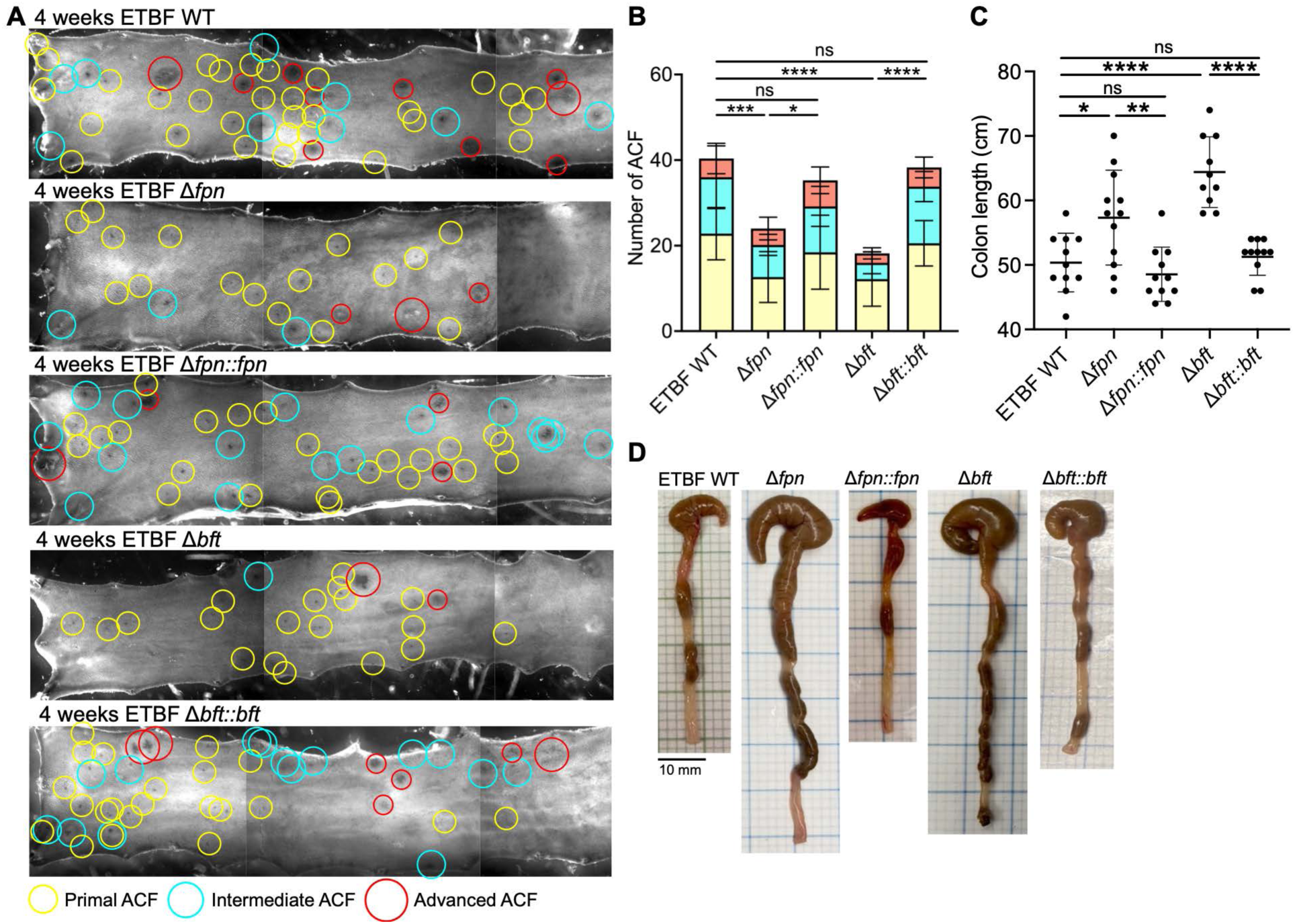
Genetic complementation restores ETBF-induced tumor-associated phenotypes in *Apc*^Min/+^ mice, related to. Fig. 3. (A) Representative methylene blue-stained distal colons from *Apc*^Min/+^ mice neonatally colonized with ETBF WT, Δ*fpn*, Δ*fpn*::*fpn*, Δ*bft*, or Δ*bft*::*bft*. ACF lesions are classified as primal (1-2 crypts, yellow), intermediate (3-6 crypts, cyan), or advanced (>7 crypts, red). (B) Quantification of primal, intermediate, and advanced ACF in the indicated colonization groups. (C) Quantification of colon length in the indicated groups. (D) Representative gross intestinal anatomy from the indicated colonization groups. n = 11, 12, 11, 10, and 11 mice per group, respectively.

**Extended Data Fig. 7.**
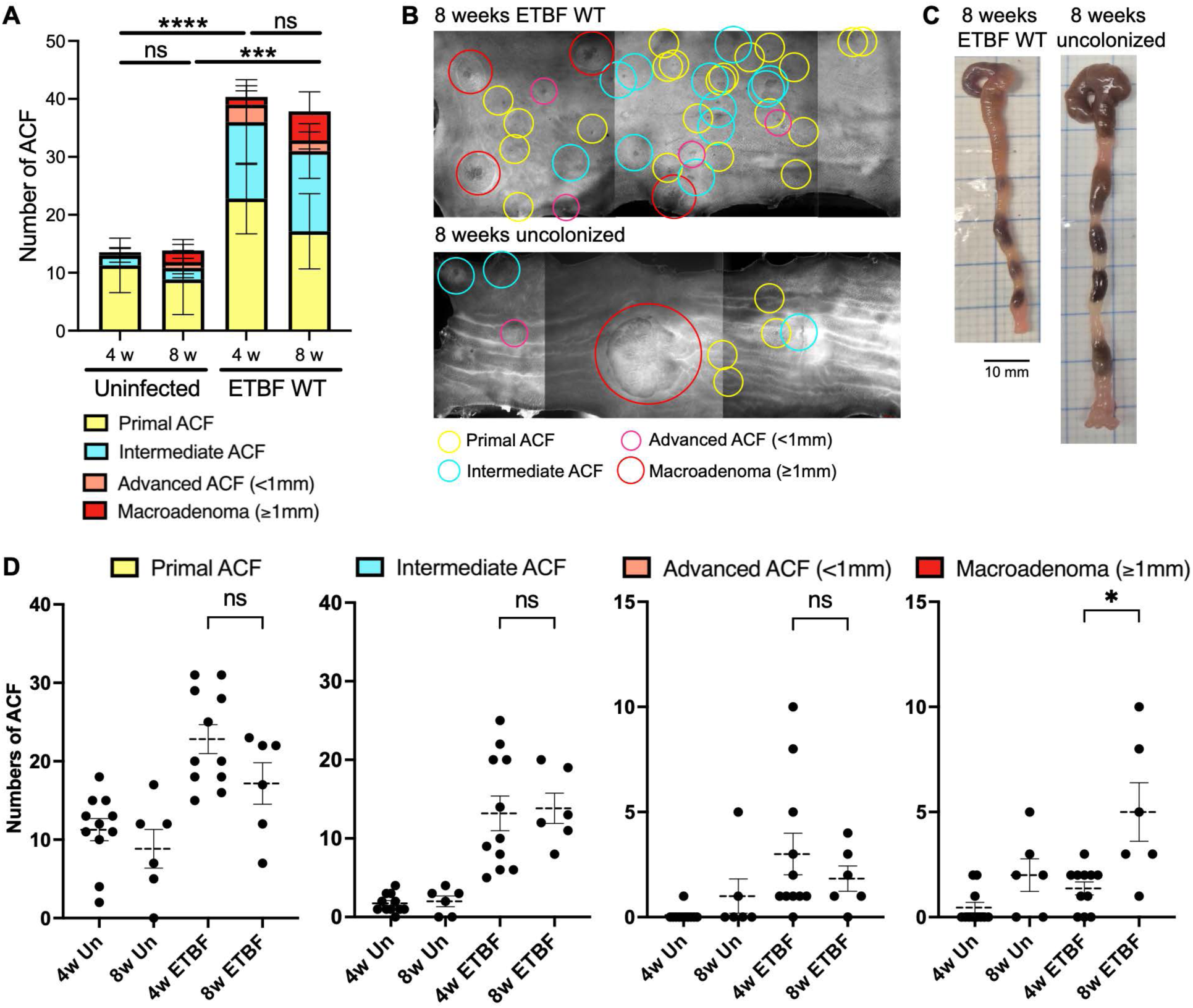
Progression of ACF lesions in *Apc*^Min/+^ mice neonatally colonized with ETBF, related to. Fig. 3. *Apc*^Min/+^ mice neonatally colonized with ETBF WT or left uncolonized were analyzed at 4 and 8 weeks of age. (A) Total ACF counts were quantified in ETBF WT-colonized and uncolonized groups at both time points. (B) Representative methylene blue-stained colons from ETBF WT-colonized and uncolonized mice at 8 weeks. (C) Representative gross colon images at 8 weeks. (D) Quantification of primal, intermediate, advanced (<1 mm), and macroadenoma (≥1 mm) lesions in each group. n = 11, 6, 11, and 6 mice for 4-week uncolonized, 8-week uncolonized, 4-week ETBF WT, and 8-week ETBF WT groups, respectively. The 4-week ETBF WT group is the same cohort shown in Extended Data Fig. 6. Data are mean ± SD. *p < 0.05, ****p < 0.0001, ns = not significant.

**Extended Data Fig. 8.**
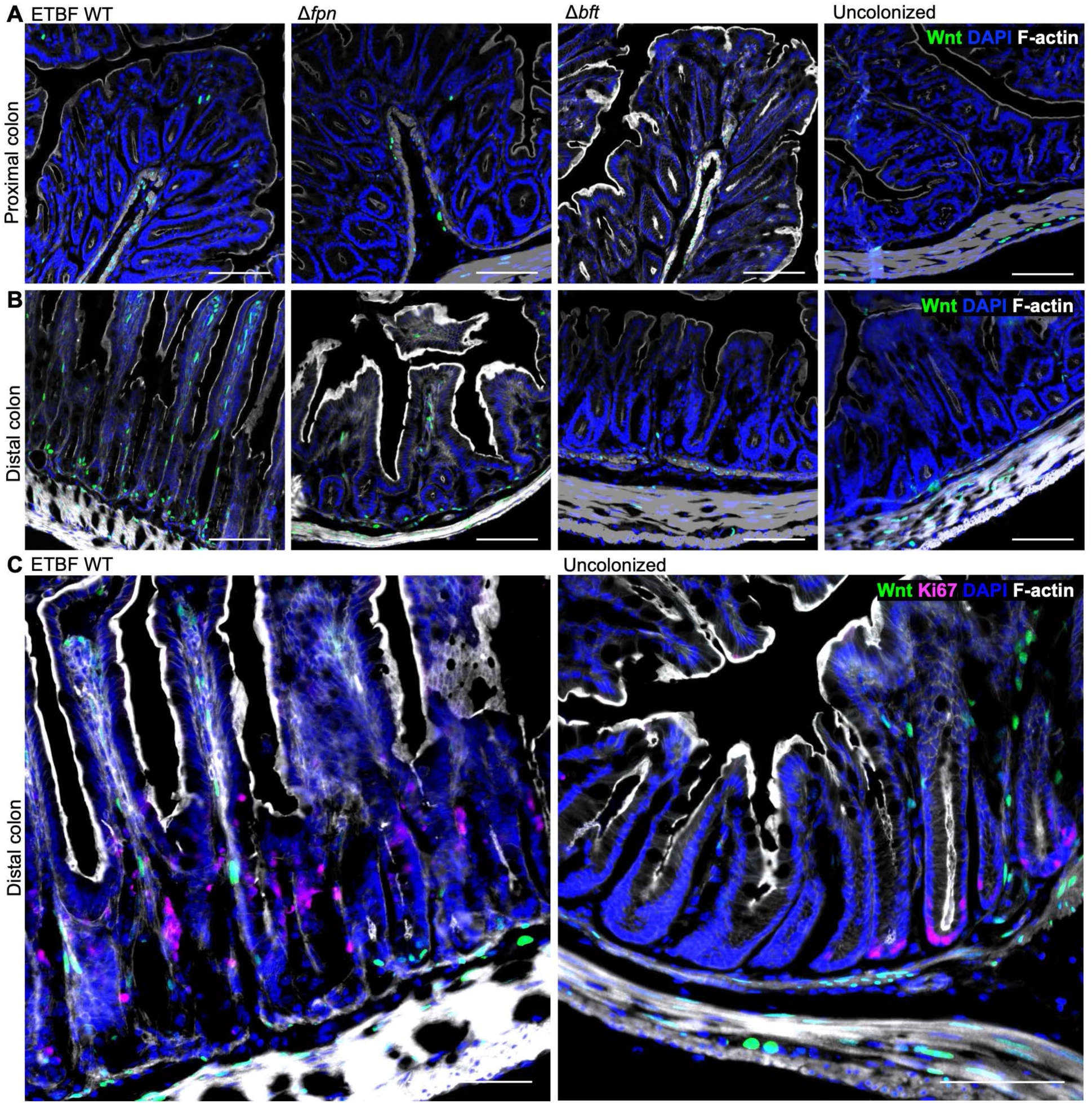
ETBF induces distal colon-specific Wnt hyperactivation and expansion of the proliferative zone, related to. Fig. 4. Wnt activity reporter mice were neonatally colonized with ETBF WT, Δ*fpn*, or Δ*bft* strains, or left uncolonized, and analyzed at 3 weeks of age. (A) Proximal and (B) distal colon sections were stained for F-actin (white) and DAPI (blue); (C) magnified distal colon sections also include Ki67 (magenta). Scale bars, 100 μm.

**Extended Data Fig. 9.**
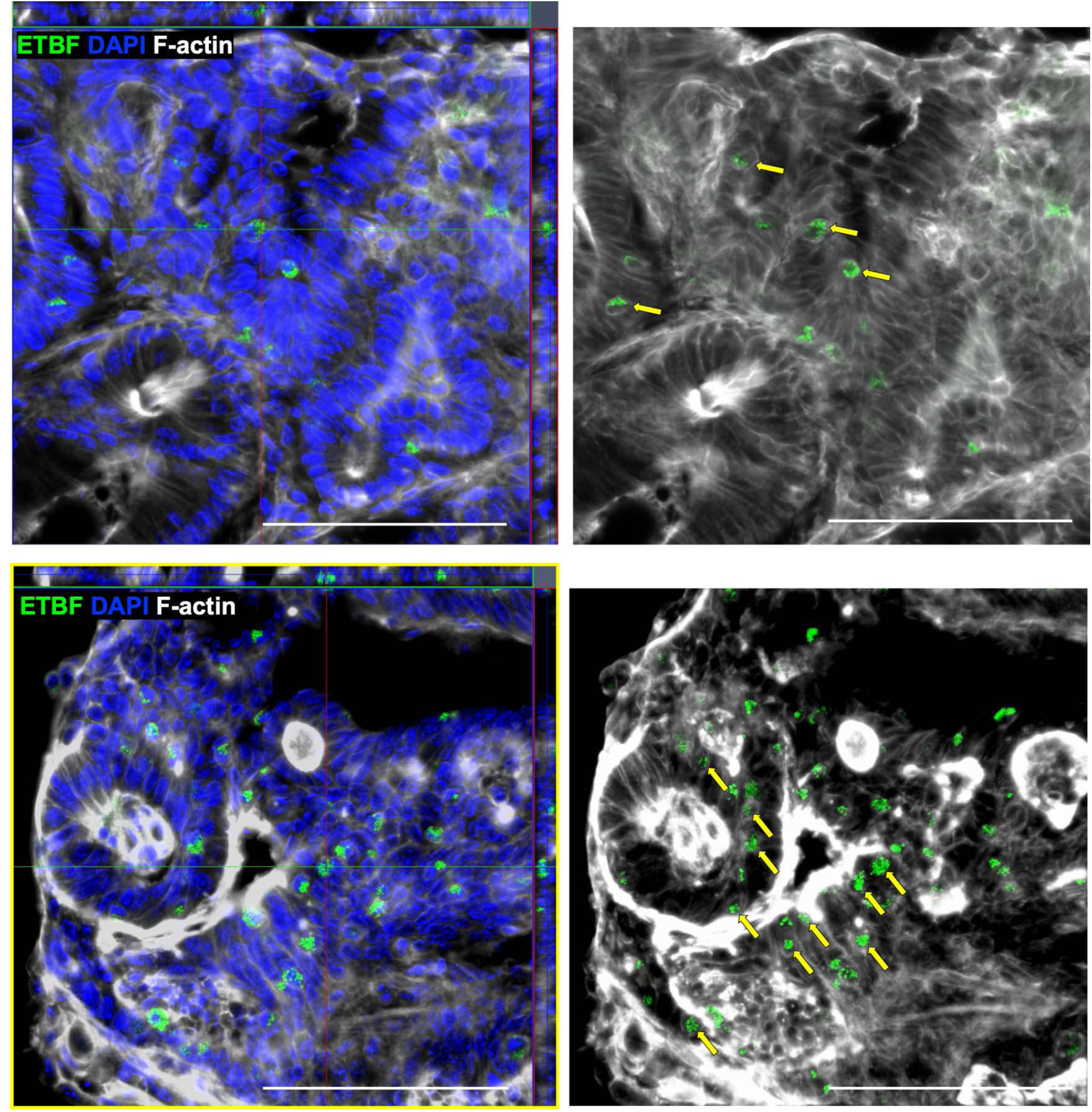
Epithelial localization of ETBF within ACF lesions following neonatal colonization, related to. Fig. 5. Representative confocal z-stack images of distal colon from 8-week-old *Apc*^Min/+^ mice neonatally colonized with ETBF WT, showing ACF lesions. ETBF (green), DAPI (blue), and F-actin (white) are shown. The top and bottom panels show independent ACF lesions. Orthogonal reconstructions (left) and maximum intensity projections (right) demonstrate localization of bacterial aggregates within the epithelial compartment (yellow arrows). Projection images are shown without DAPI to improve visualization of bacterial localization. Scale bars, 100 μm.

**Extended Data Fig. 10.**
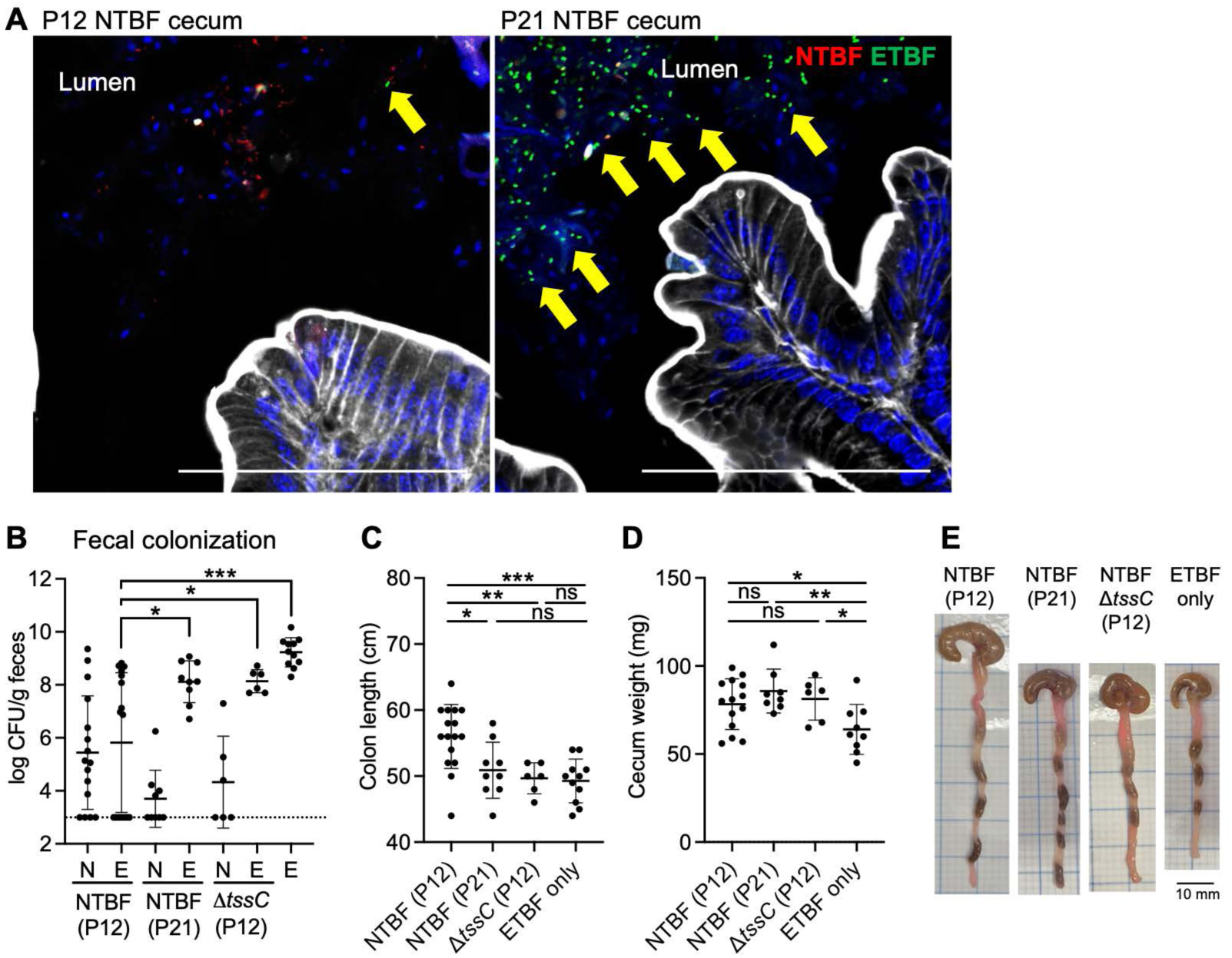
Timing and T6SS dependence of NTBF-mediated protection against ETBF colonization and pathology, related to. Fig. 6. *Apc*^Min/+^ pups born to ETBF-colonized dams were gavaged with NTBF at P12 or P21, NTBF Δ*tssC* at P12, or left untreated (ETBF only), and analyzed at 4 weeks of age. (A) Representative confocal images showing luminal ETBF-sfGFP (green, yellow arrows) and NTBF-mCherry (red). (B) Fecal bacterial burden was assessed (N = NTBF, E = ETBF). (C) Cecal length, (D) cecal weight, and (E) representative gross colon images. Scale bars, 100 μm (A). Two independent replicates were performed; n = 14 (P12 NTBF), n = 8 (P21 NTBF), and n = 6 (P12 Δ*tssC*), and n = 11 (ETBF only). *p < 0.05, **p < 0.01, ***p < 0.001, ns = not significant.

## Notes

### Competing Interest Statement

The authors have declared no competing interest.

