## Supplementary Information for "Early-life colonization by enterotoxigenic *Bacteroides fragilis* remodels gut epithelial stem cell states to drive colorectal cancer susceptibility"

### Early-life bacterial toxin exposure remodels gut epithelial stem cell fate to drive cancer susceptibility

**The PDF file includes:**

Supplementary Figs. 1-3

Supplementary Tables 1-3

**Other Supplementary Materials for this manuscript include the following:**

Supplementary Movies 1-2

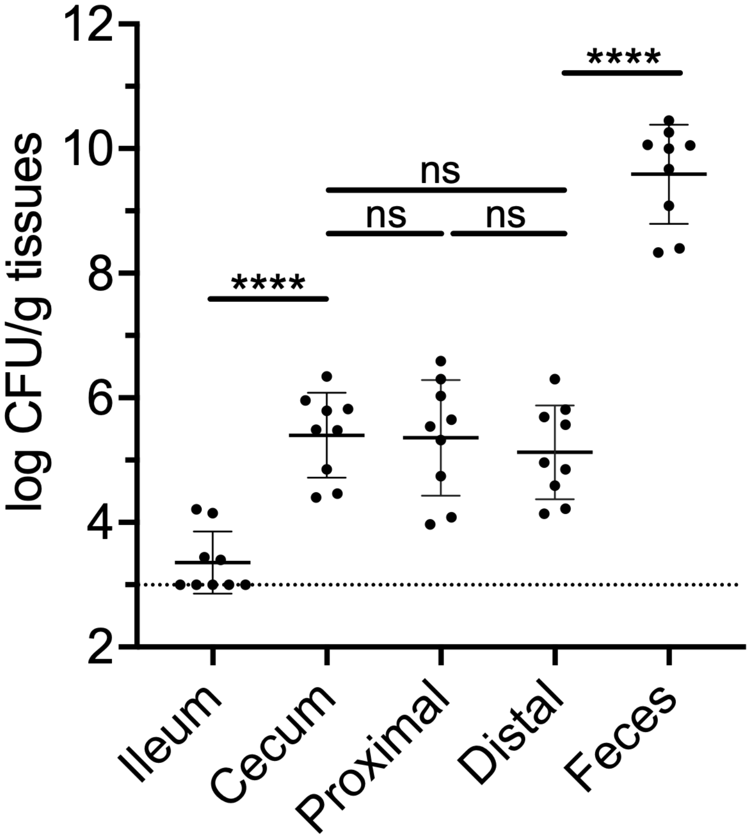


**Supplementary Fig. 1. Quantification of ETBF tissue colonization across intestinal regions, compared with fecal titers.** Intestinal tissues were extensively washed with PBS (10–15 times per cycle, 3 cycles total) to remove luminal bacteria, including those within the mucus layer, prior to homogenization. Tissue homogenates were serially diluted and plated for colony-forming unit (CFU) enumeration. Data represent CFU per tissue from the indicated intestinal locations. ****p < 0.0001, ns = not significant.

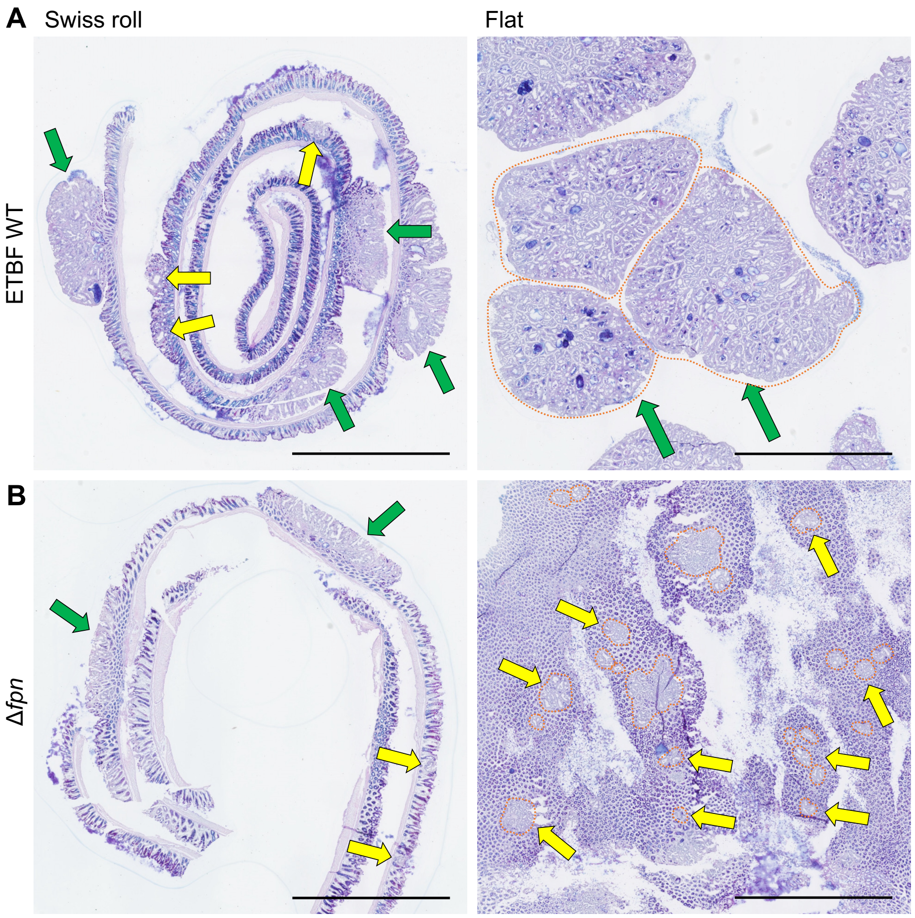


**Supplementary Fig. 2. Visualization of precancerous lesions in distal colon tissue.** Distal colons from 8-week-old *Apc*$^{\mathrm{Min/+}}$ mice neonatally colonized with ETBF WT or Δ*fpn* strains were analyzed histologically and by confocal microscopy. (A-B) AB/PAS-stained Swiss-roll (left) and flat-mounted (right) sections show aberrant crypt foci (ACF; yellow arrows) and early polyps (green arrows) in colons colonized with (A) ETBF WT or (B) Δ*fpn*.

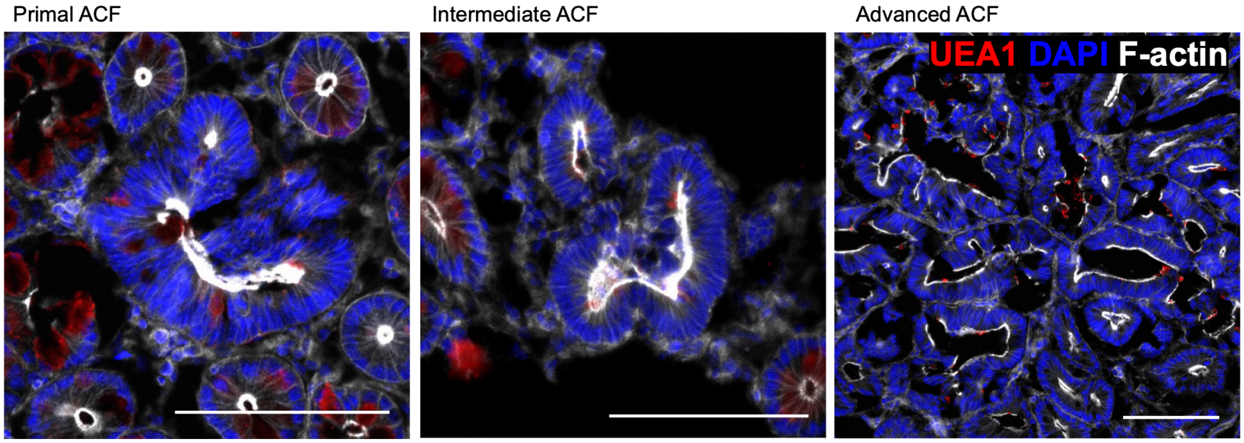


**Supplementary Fig. 3. UEA1 staining confirms reduced goblet cell-associated mucin signal in ETBF-induced ACF lesions, related to Fig. 3.** Representative confocal images of flat-mounted distal colon tissue from ETBF-colonized *Apc*$^{Min/+}$ mice stained with UEA1 (red), DAPI (blue), and F-actin (white), showing primal, intermediate, and advanced ACF lesions. ACF lesions show reduced UEA1 signal compared to adjacent normal crypts. Scale bars, 100 µm.

**Supplementary Table 1.**

Bacterial strains used in this study.

| Strain | Sources | Identifier | Variants |
|---|---|---|---|
| *B. fragilis* (ETBF) | ATCC;<br>This study;<br>Hill et al., 2024 (*30*);<br>Hecht et al., 2016 (*31*) | ATCC 43859 | GFP Tet$^{R}$Clin$^{R}$<br>Δ*bft* GFP Tet$^{R}$Clin$^{R}$<br>Δ*fpn* GFP Tet$^{R}$Clin$^{R}$ |
| *B. fragilis* (NTBF) | ATCC;<br>Hill et al., 2024 (*30*);<br>Hecht et al., 2016 (*31*) | TM4000/638R | RFP Rif$^{R}$Clin$^{R}$<br>Δ*tssC* RFP Rif$^{R}$Clin$^{R}$ |

**Supplementary Table 2.**

Mouse strains and genotypes used in this study. R26-LSL-H2B-mCherry mice were crossed with Atoh1-Cre mice to generate Atoh1 (R26-LSL-H2B-mCherry; Atoh1-Cre) reporter mice used in this study.

| Mouse line | Description / Genotype | Experimental use | Source / Reference |
|---|---|---|---|
| C57Bl/6J | Inbred wild-type background strain | Baseline control and background strain; used for breeding and neonatal colonization studies | The Jackson Laboratory (000664) |
| Lgr5-EGFP-IRES-CreERT2 | Knock-in Lgr5-EGFP reporter | Lineage tracing and quantification of stem-cell expansion | JAX (008875) |
| TCF/Lef:H2B-GFP | Nuclear GFP reporter of Wnt co-factor TCF/Lef | Detection of Wnt activation | JAX (013752) |
| Atoh1-Cre | Cre recombinase knocked-in at the *Atoh1* locus | Secretory-lineage-specific recombination for epithelial tracing | JAX (011104) |
| R26-LSL-H2B-mCherry | Rosa26-loxP-STOP-loxP nuclear mCherry reporter | Visualization of Cre-dependent labeling of epithelial lineages | JAX (023139) |

| | | | |
|---|---|---|---|
| *Apc*$^{\text{Min/+}}$ | Germline truncating mutation in *Apc* tumor-suppressor gene | Model of spontaneous intestinal adenoma and ETBF-driven tumorigenesis | JAX (002020) |

**Supplementary Table 3.**

Antibodies and probes used for immunofluorescence.

| Target | Antibody / probe | Dilution | Source |
|---|---|---|---|
| F-actin | Phalloidin (Alexa Fluor™ 555) | 1:250 | Invitrogen A34055 |
| F-actin | Phalloidin (Alexa Fluor™ 647) | 1:250 | Invitrogen A22287 |
| Nucleus | ProLong™ Gold Antifade Mountant with NucBlue™ (DAPI) | Ready-to-use | Thermo Fisher Scientific P36931 |
| Goblet cells/ mucin | DyLight™ 649–conjugated Ulex europaeus agglutinin I (lectin) | 1:200 | Vector Laboratories DL-1068-1 |
| Ki-67 | Rabbit anti-Ki-67 | 1:250 | Abcam ab15580 |
| Secondary antibody | Alexa Fluor™ 647–conjugated goat anti-rabbit IgG | 1:1000 | Invitrogen A21244 |

**Supplementary Movie 1.**

Three-dimensional confocal Z-stacks reconstructed with Volocity software show sfGFP-expressing ETBF aggregates within distal colonic ACF lesions of 8-week-old *Apc*$^{Min/+}$ mice following neonatal ETBF colonization. DAPI (blue) and F-actin (white) outline crypt architecture, highlighting the spatial association of ETBF with early neoplastic crypts. Imaged at 20× magnification.

**Supplementary Movie 2.**

Same as Movie S1, imaged at 40× magnification.
